# SMAD4 suppresses colitis-associated carcinoma through inhibition of CCL20/CCR6-mediated inflammation

**DOI:** 10.1101/2022.05.25.492945

**Authors:** David N. Hanna, Paula Marincola Smith, Sergey V. Novitskiy, M. Kay Washington, Jinghuan Zi, Connie J. Weaver, Jalal A. Hamaamen, Keeli B. Lewis, Jing Zhu, Jing Yang, Qi Liu, R. Daniel Beauchamp, Anna L. Means

## Abstract

**Background & Aims:** Chronic inflammation in the colon is a predisposing factor for colon cancer. We previously reported that colon epithelial cell silencing of *Smad4*, the central downstream mediator of TGFβ family signaling, increased epithelial expression of inflammatory genes, including the chemokine CCL20, and increased susceptibility to colitis-associated cancer. Here, we examine the role of the chemokine/receptor pair CCL20/CCR6 in mediating colitis-associated colon carcinogenesis induced by SMAD4 loss.

**Methods:** Mice with conditional, epithelial-specific *Smad4* loss with and without germline deletion of the *Ccr6* gene were subjected to three rounds of dextran sodium sulfate and followed for up to 3 months. Tumors were quantified histologically, and immune cell populations were analyzed by flow cytometry and immunostaining.

**Results:** In humans, *SMAD4* expression inversely correlated with *CCL20* expression. *Smad4* loss in mouse colon epithelium led to enlarged gut-associated lymphoid tissues and recruitment of specific immune cell subsets to the mouse colon epithelium and underlying stroma, particularly Treg, T_h_17, and dendritic cells. Loss of CCR6 abrogated these immune responses and significantly reduced the incidence of colitis-associated tumors observed with loss of SMAD4 alone.

**Conclusions:** Regulation of mucosal inflammation is a critical role of SMAD4 signaling within the colon epithelium and central to its tumor suppressor function in the colon. A key downstream node in this regulation is suppression of CCL20 signaling by the epithelium to CCR6 in immune cells. Loss of SMAD4 in the colon epithelium increases CCL20 expression and chemoattraction of CCR6+ immune cells, contributing to greater susceptibility to colon cancer.

## Introduction

Chronic inflammation is a predisposing factor for many cancers, including colorectal cancer (CRC), which is the third most common cancer worldwide ^1, 2^. Inflammatory bowel disease (IBD) patients exemplify this relationship, as they demonstrate a markedly elevated risk of developing CRC, often early in life ^3^.

Transforming growth factor β (TGFβ) family signaling functions in a critical tumor suppressor role, particularly within the alimentary tract. In the canonical pathway, TGFβ signaling is dependent on SMAD4 as the pathway’s common mediator ^4^. Previous work from our lab demonstrated that TGFβ signaling via SMAD4 blocks or represses the effects of pro-inflammatory cytokines in colonic epithelium ^5^. Additionally, we found that conditional knockout of the *Smad4* gene in murine intestinal epithelial cells led to a striking increase in epithelial inflammatory signaling, stromal leukocyte infiltration, and profoundly increased susceptibility to colitis-associated cancer (CAC) development compared to SMAD4+ control mice ^5^.

We previously identified that expression of C-C motif chemokine ligand 20 (CCL20) is markedly increased in colon epithelial cells *in vivo* and *in vitro* after loss of SMAD4 or inhibition of TGFβ receptors ^5^. CCL20 acts via its sole receptor, C-C motif chemokine receptor (CCR6), and is expressed on multiple immune cell subtypes including T_h_17 cells, Tregs, B cells, multiple dendritic cell (DC) subsets, and others ^6, 7^. In humans, CCL20 is upregulated in both ulcerative colitis and Crohn’s disease, and is elevated in sporadic colorectal adenomas and in CRC, suggesting that it may influence cancer susceptibility ^8-10^.

Here we demonstrate that *SMAD4* mRNA expression is negatively correlated with *CCL20* mRNA expression in human IBD and CRC specimens and that human ulcerative colitis-associated cancers with SMAD4 loss have increased immune cell infiltration. Furthermore, we found that conditional colon epithelial cell *Smad4* gene loss in mice leads to increased expression of inflammation-related genes within the colonic sub-epithelial stromal tissues, including *Ccr6*, and an increase in gut-associated lymphoid tissue (GALT) area. Following dextran sodium sulfate (DSS) -induced colitis, specific immune populations were increased by SMAD4 loss, including stromal Tregs and stromal and epithelial-associated CD4+ IL17a+ T cells (T_h_17) and DC subtypes. Notably, these cells are all known to be regulated by CCR6 activity and loss of *Ccr6* abrogated much of the phenotype that was induced by epithelial *Smad4* loss alone. Furthermore, we demonstrate that CAC development due to epithelial SMAD4 loss is dependent on an intact CCL20/CCR6 axis.

## Materials and Methods

### In silico Analysis of Microarray Data and The Cancer Genome Atlas (TCGA)

Gene expression in IBD patients and healthy controls was analyzed from published transcriptomic data generated from endoscopic biopsy samples and analyzed by Microarray (GSE75214) ^11^ as we have previously published ^12^. To determine the relationship between *SMAD4* and *CCL20* expression in sporadic colon cancer specimens, rectal cancer specimens, and healthy representative controls, the TCGA database (https://www.cancer.gov/tcga) was analyzed utilizing the Firehose web browser from the Broad Institute (https://gdac.broadinstitute.org/).

### Multiplex Immunofluorescent (mxIF) staining of Human Tissues

Tissue microarrays (TMAs) were created from de-identified ulcerative colitis-associated cancers (UCACs) and SMAD4 status was previously determined by immunohistochemistry ^5^. Sequential staining (Supplementary Table S3), single-cell segmentation, and cell quantification was performed by the Vanderbilt University Medical Center (VUMC) Digital Histology Shared Resource.

### Mouse Model

Animal work was performed with approval from the Vanderbilt University Institutional Animal Care and Use Committee and followed ARRIVE guidelines. We previously published that tamoxifen-treated *Lrig1*^*CreERT*^*Smad4*^*fl/fl*^ (termed *Smad4*^*ΔLrig1*^) mice have greater than 90% loss of SMAD4 in their colon epithelium ^5^. SMAD4-expressing control mice (*Smad4*^*fl/fl*^ mice with no Cre alleles) treated with tamoxifen are referred to as SMAD4+ or control mice for simplicity.

For animal experiments involving mice with or without *Ccr6* expression, mice with *Lrig1*^*CreERT2*^ and *Smad4*^*fl/fl*^ alleles were bred with mice from Jackson Laboratory (Bar Harbor, ME) that have *GFP* knocked into the *Ccr6* coding region such that cells transcribing from the *Ccr6* promoter can be tracked by GFP protein expression in heterozygous *Ccr6*^*GFP/+*^ mice and in *Ccr6*^*GFP/GFP*^ null mice ^13^.

Experimental colitis was induced using 2% DSS (MP Biomedicals, Santa Ana, CA) in drinking water for 5 days, followed by 5 days of recovery, in 3 consecutive cycles. Controls were sibling littermates and cage mates. After tamoxifen treatment, bedding was mixed among cages within an experiment once per week.

### RNA-Sequencing

For bulk RNAseq, three *Smad4*^*ΔLrig1*^ and three SMAD4+ control mice were dissected one month after administration of tamoxifen. Colons were stripped of crypts as published ^5^. Stroma was scraped from underlying muscle and RNAseq was performed by the Vanderbilt Technologies for Advanced Genomics (VANTAGE) core facility. The data files are publicly available on the National Institute of Health Gene Expression Omnibus database (GSE189667) _14, 15_.

### *RNAScope ®* In Situ *Hybridization*

*In situ* hybridization by RNAscope® technique was performed according to the manufacturer’s instructions on 5μm thick sections from five *Smad4*^*ΔLrig1*^ and five SMAD4+ control mice [Advanced Cell Diagnostics (ACD), Newark, CA, USA]. To quantify differential *Ccl20* expression, specific colorimetric dots corresponding with mRNA copies were counted in each positive cell. To quantify cells expressing *Ccr6*, the number of positive cells were counted among the distalmost 100 crypts.

### Mouse Histology and Immunostaining

Mouse tissues were processed as described ^16^. For analysis of GALTs, three 5μm sections were cut and examined per mouse, each 1000μm apart. Sections were stained by hematoxylin and eosin (H&E) and examined for frequency and cross-sectional GALT area normalized to total tissue area present. For immunohistochemical (IHC) or immunofluorescent (IF) analyses of cell types associated with the epithelium, cells were counted in or immediately adjacent to the distal-most 100 crypts, respectively. For quantification of cell types within GALTs, cell counts were normalized to GALT cross-sectional area for the distalmost five GALTs. For evaluation of tumor development and inflammation, a collaborating expert pathologist (MKW), who was blinded to mouse genotype, examined six sections per mouse that were each 200μm apart for presence and number of invasive tumors as well as histologic scoring of colitis as previously described ^17^.

### Flow Cytometry

Six SMAD4+ *Ccr6*^*GFP/+*^, five SMAD4+ *Ccr6*^*GFP/GFP*^, five *Smad4*^*ΔLrig1*^*Ccr6*^*GFP/+*^, and four *Smad4*^*ΔLrig1*^*Ccr6* ^*GFP/GFP*^ mice were given three rounds of DSS in drinking water as described above. One month after conclusion of DSS treatment, the mice were euthanized, and single cells were isolated from colon epithelium and stroma ^5, 18^. Single cell suspensions from both the epithelium and stroma were aliquoted for staining in two parallel panels as displayed in Supplementary Figure 1A (BioLegend, San Diego, CA). Since fluorescence from GFP produced from the *Ccr6* promoter could not be detected directly, we used an anti-GFP antibody to detect cells that express the *Ccr6* promoter, even in *Ccr6*^*GFP/GFP*^ mice that do not produce CCR6 protein (Supplementary Figure 1B). Data were collected on a 5-laser BD Biosciences LSRII Flow Cytometer (San Jose, CA) in the VUMC Flow Cytometry Shared Resource Core.

### Statistics

*In silico* analysis of gene expression data in human biopsy samples were compared using the non-parametric Kruskal-Wallis test with post hoc Welch’s *t* test (IBD specimens), non-parametric Mann-Whitney test (TCGA specimens), or non-parametric Spearman’s correlation (IBD and TCGA specimens), as appropriate. Mann-Whitney U test was used to compare GALT area, immunostaining cell quantities, colitis inflammation scores, mean number of tumors between groups *in vivo* following DSS treatment, and human CAC mxIF immunophenotyping. Fisher’s Exact test was used to compare proportion of animals in each group that developed tumors following DSS treatment. Flow cytometry results were analyzed by multiple 2-tailed T test with Holm-Sidak multiple test correction. Statistical analyses were performed using GraphPad Prism 9 Software (San Diego, CA). Throughout the manuscript, statistical significance is designated as: ns (*p* ≥ .05), * (*P* < .05), and ** (*P* < .001).

### Study Approval

All animal work was performed after approval by the Vanderbilt Institutional Animal Care and Use Committee. Acquired human tissue was de-identified and used with approval from the Vanderbilt Institutional Review Board.

## Results

### SMAD4 *expression is negatively correlated with* CCL20 *expression in human IBD and CRC specimens*

Our prior work demonstrated that loss of *Smad4* in colon epithelial cells was associated with significant upregulation of multiple inflammation-related genes, including *Ccl20* ^5^. Additionally, we demonstrated that *SMAD4* expression was significantly decreased in active ulcerative colitis (Uca) and active Crohn’s Disease (CDa) specimens compared to healthy colon (HC) biopsy specimens analyzed by DNA Microarray (GSE75214 ^11^) ^11, 12^. Analysis of the same database revealed that *CCL20* expression was significantly increased in UCa, UCi (inactive ulcerative colitis), and CDa specimens compared to HC and that a significant inverse correlation exists between *SMAD4* and *CCL20* expression in human IBD specimens (r^2^ = -0.30, *P =* .001; Figure 1A).

**Figure 1.**
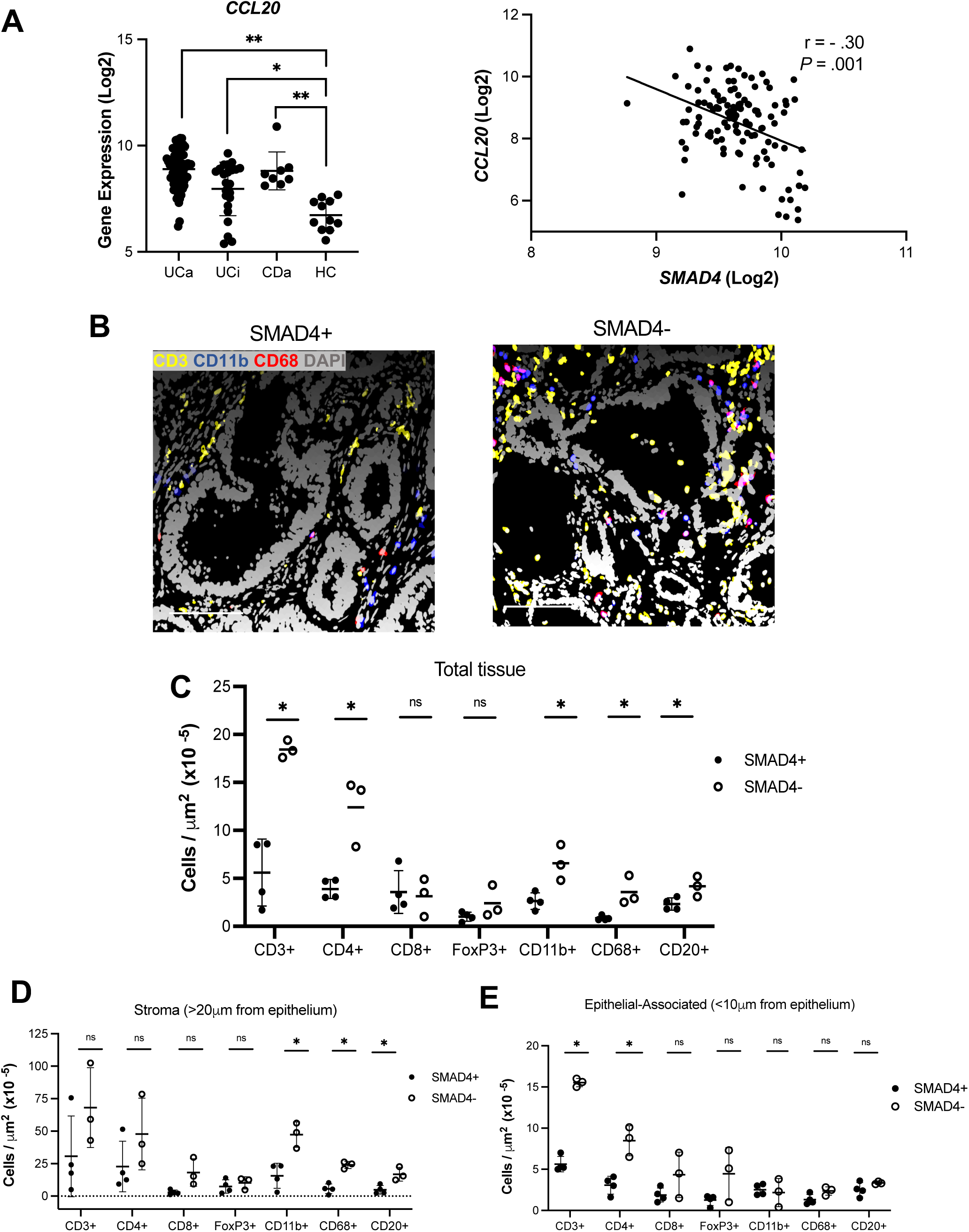
*SMAD4* expression is negatively correlated with *CCL20* gene expression in human IBD and sporadic colon and rectal cancer and is associated increased immune cell recruitment in human ulcerative colitis-associated cancer specimens. **(A)** *In silico* analysis of human microarray data from IBD colonoscopic biopsies (UCa: active ulcerative colitis; UCi: inactive ulcerative colitis; CDa: active Crohn’s disease; HC: healthy control). **(B)** MxIF of human UCACs stained for indicated markers. **(C)** Quantification of immune cell types relative to total tissue area. **(D)** Quantification of stromal and **(E)** epithelial-associated immune cells relative to their respective tissue area (CD3 = yellow, CD11b = blue, CD68 = red, DAPI = gray; scale bars: 100μm; data points represent mean per tumor sample if multiple cores exist; (\**P* < .05, ** *P* < .001)

We have previously shown that *SMAD4* expression is significantly decreased in colon cancer (CC) specimens relative to healthy colon (HCc) specimens by analysis of the TCGA COAD colorectal cancer database ^12^. Analysis of the same database revealed that *CCL20* was significantly increased in CC compared to HCc specimens, and there was a significant negative correlation between *SMAD4* and *CCL20* expression in these tissues (r^2^ = -0.24, *P* < .001; Supplementary Figure 2A). We observed a similar relationship in rectal cancer (RC) by TCGA analysis (r^2^ = -0.299, *P=*.001; Supplementary Figure 2B). Thus, the relationship that we previously observed between *Smad4* and *Ccl20* expression in mouse colons is reflected in human IBD and sporadic CRC.

To investigate whether decreased SMAD4 protein expression is associated with altered immune cell infiltration in colitis-associated cancers, we performed multiplexed immunofluorescence (mxIF) on a tissue microarray containing three tissue cores from three SMAD4- and seven cores from four SMAD4+ human UCAC specimens. We found that SMAD4-UCACs exhibited an increase in overall T cell infiltration (particularly CD4+ T cells), as well as CD11b+ monocytes, CD68+ macrophages, and CD20+ B cells normalized to total tissue area when compared to SMAD4+ human UCACs (Figure 1B-C). Examining specific tissue compartments, we found that the stroma surrounding, but not in contact with the epithelium, in SMAD4-UCACs exhibited a greater number of CD11b+ monocytes, CD68+ macrophages, and CD20+ B cells compared to SMAD4+ tumors coincident with significantly increased epithelium-associated CD3+ and CD4+ T cells (Figure 1D-E). Thus, loss of SMAD4 alters key immune regulators both in close association to epithelium, as well as throughout the tumor microenvironment. In consideration of these findings in human UCACs, we further explored the effect of colon epithelial cell *Smad4* loss in the mouse colon and in the setting of colitis in mice.

### *Intestinal Epithelial* Smad4 *loss results in increased expression of inflammatory signaling genes in mouse colonic stroma, including* Ccr6

To examine how increased epithelial inflammatory signaling in the setting of *Smad4* loss impacts stromal composition under homeostatic conditions, we performed bulk RNAseq on the isolated sub-epithelial stroma from three *Smad4*^*ΔLrig1*^ and three SMAD4*+* control mice. 95 genes were upregulated at least 2.5-fold (≥1.32 Log_2_ Fold Change) with a false discovery rate of <0.05 (Supplementary Figure 3A, Supplementary Table 1). Of those 95 genes, 51 are known to be expressed in immune cells or have immune-related functions (Supplementary Table 2). Ingenuity pathway analysis demonstrated that the top 15 most significantly altered signaling pathways are immune-related, including pathways related to Th1/Th2 signaling, B cell development, leukocyte extravasation, and DC maturation, among others (Supplementary Figure 3B). ImmuCC analysis of the RNA-seq data predicted a relative increase in B cells and CD4+ T cells and a simultaneous relative decrease in CD8+ T cells and monocytes in the colonic stroma of *Smad4*^*ΔLrig1*^ mice relative to SMAD4+ controls (Supplementary Figure 3C). Among the upregulated stromal genes detected were chemokine receptors, including *Ccr7, Cxcr4, Cxcr5, and Ccr6*, the latter of which encodes CCL20’s receptor, CCR6 (Supplementary Table 1-2). RNAScope *In situ* hybridization confirmed that *Smad4*^*ΔLrig1*^ mice exhibited significantly more *Ccr6+* stromal cells relative to SMAD4+ mice (265.2 vs 94.8 cells /100 crypts, *P <*.001; Supplementary Figure 3D). These findings, together with our prior findings that loss of epithelial SMAD4 increases CCL20 expression^5^, suggest that *Smad4* loss in colon epithelial cells results in a pro-inflammatory stromal response involving the CCL20/CCR6 signaling axis.

### Smad4 *loss results in altered GALT phenotype, which is dependent on an intact CCL20/CCR6 axis*

GALTs have a critical role in mucosal immunity, serving as major sites of T and B cell recruitment and activation, antigen sampling, and induction of local immune responses ^19^. CCL20 is constitutively produced by follicle-associated epithelial (FAE) cells that cover the dome of GALTs where it is thought to contribute to recruitment of CCR6+ leukocytes, particularly B cells, T cells, and DCs, to GALTs ^20, 21^. RNAScope *In situ* hybridization demonstrated that *Smad4*^*ΔLrig1*^ mice exhibited significantly increased *Ccl20* expression in the follicle-associated epithelial cells immediately adjacent to GALTS compared to SMAD4+ mice (35.3 vs 12.2 mRNA copies/ positive cell, *P <*.001; Supplementary Figure 4A). To determine if the increased *Ccl20* expression occurring with loss of colon epithelial *Smad4* is associated with alterations in GALT morphology, we examined the colons of five *Smad4*^*ΔLrig1*^ and five SMAD4+ control mice for frequency and size of GALTs under homeostatic conditions. While *Smad4*^*ΔLrig1*^ mice did not exhibit significantly more GALTs per colon than SMAD4+ control mice (17.2 vs 14.7, *P =*.35), GALTs in *Smad4*^*ΔLrig1*^ mice were significantly larger than GALTs in SMAD4+ control mice (6.13×10^4^ μm^2^ vs 4.80 ×10^4^ μm^2^, *P =* .005; Figure 2A).

**Figure 2.**
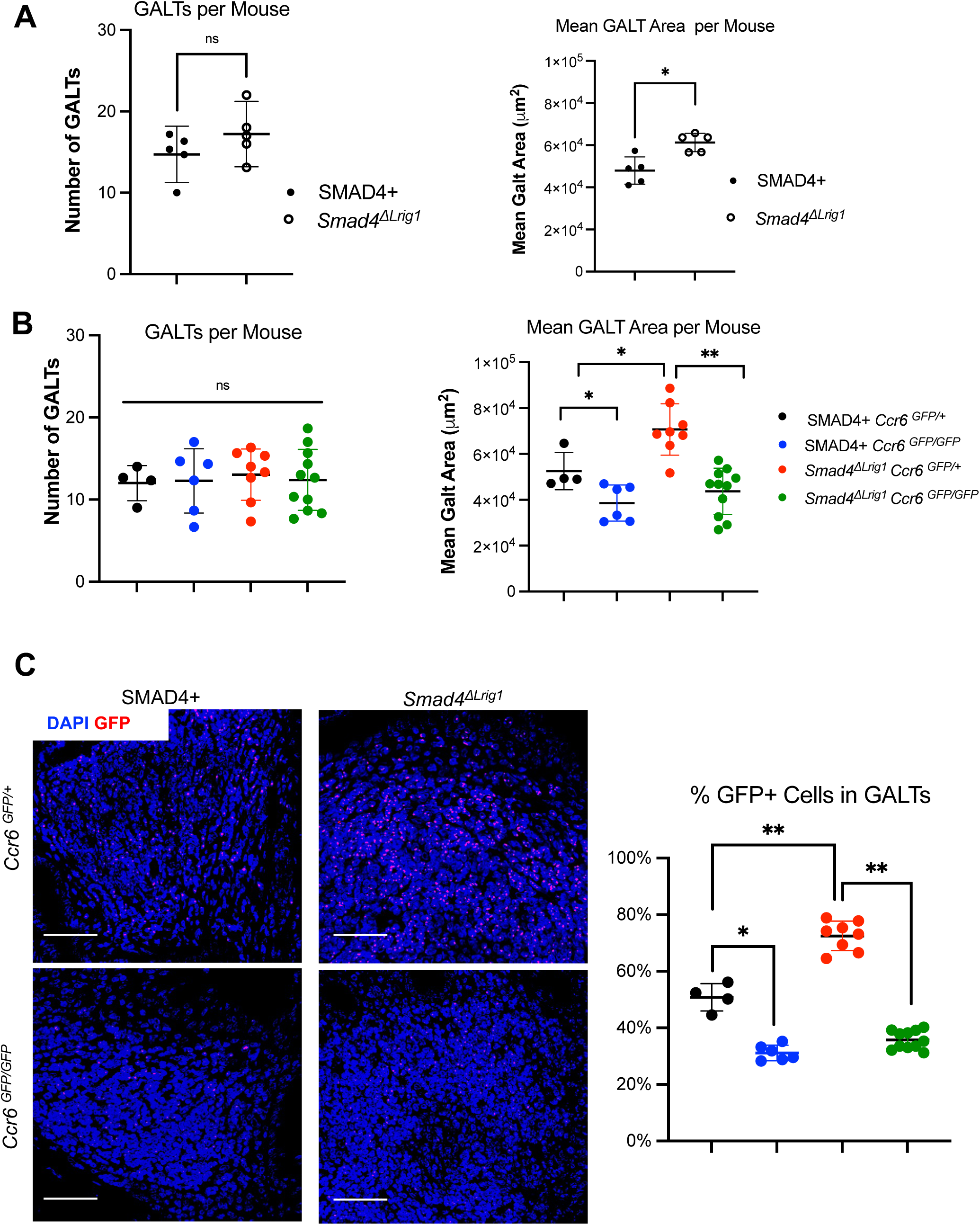
*Smad4* loss results in increased GALT area only when CCR6 is expressed. **(A)** Quantification of GALT number and cross-sectional area from *Smad4*^*ΔLrig1*^ and SMAD4+ control mice under homeostatic conditions, **(B)** and in mice with or without expression of *Smad4* or *Ccr6*, 3 months after completion of DSS-induced colitis. **(C)** Immunofluorescent staining and quantification for GFP (red) in GALTS from indicated genotypes. Data points represent mean quantification per mouse (DAPI: blue, scale bars: 100μm; (\**P* < .05, ** *P* < .001).

To determine if the altered GALT morphology persisted with inflammation and was dependent on CCR6 expression, the same analysis was performed in SMAD4+ *Ccr6* ^*GFP/+*^, SMAD4+ *Ccr6*^*GFP/GFP*^, *Smad4*^*ΔLrig1*^ *Ccr6*^*GFP/+*^, and *Smad4*^*ΔLrig1*^ *Ccr6* ^*GFP/GFP*^ mice 3 months after completion of DSS-induced colitis. This analysis once again demonstrated that there were no significant differences in the average number of GALTs observed across mice from all four genotypes (*P =* .95; Figure 2B). However, loss of *Ccr6* decreased GALT size irrespective of *Smad4* status, with SMAD4+ *Ccr6* ^*GFP/+*^ mice having significantly larger GALTs than SMAD4+ *Ccr6*^*GFP/GFP*^ mice (5.2 ×10^4^ μm^2^ vs 3.7 ×10^4^ μm^2^, *P =* .02) and *Smad4*^*ΔLrig1*^ *Ccr6*^*GFP/+*^ mice having larger GALTs than *Smad4*^*ΔLrig1*^ *Ccr6* ^*GFP/GFP*^ mice (7.3 ×10^4^ μm^2^ vs 4.4 ×10^4^ μm^2^, *P <* .001; Figure 2B, Supplementary Figure 4B). These findings are consistent with previous reports that CCR6 serves a critical role in GALT development and immune cell trafficking in the mouse colon ^6, 22^. On the other hand, loss of *Smad4* increased GALT size when *Ccr6* was expressed, with *Smad4*^*ΔLrig1*^ *Ccr6*^*GFP/+*^ mice having larger GALTs than SMAD4+ *Ccr6* ^*GFP/+*^ mice (7.3 ×10^4^ μm^2^ vs 5.2 ×10^4^ μm^2^, *P =* .007; Figure 2B) and this gain was blocked by loss of *Ccr6*.

IF staining for GFP as a marker of *Ccr6* promoter activity revealed that an expansion of the CCR6+ cell population contributed to the larger GALT size in the context of SMAD4 loss. GALTs in *Smad4*^*ΔLrig1*^ *Ccr6*^*GFP/+*^ mice consisted of a significantly higher proportion of GFP+ cells compared to SMAD4^*+*^*Ccr6*^*GFP/+*^ mice (72.9% vs 50.9%, *P* < .001; Figure 2C) and this increase was offset by additional loss of *Ccr6* in *Smad4*^*ΔLrig1*^ *Ccr6*^*GFP/GFP*^ mice (72.9% vs 35.8%, *P* < .001). Thus, loss of epithelial SMAD4 expression leads to larger GALTs with increased proportions of CCR6+ immune cells that is dependent on *Ccr6* expression. To understand whether SMAD4 signaling alters the distribution or quantity of different immune cell types within the GALTs, IF staining was performed on tissue sections from five *Smad4*^*ΔLrig1*^*Ccr6*^*GFP/+*^ and five SMAD4+ *Ccr6*^*GFP/+*^ mice under homeostatic conditions to localize B cells, T cells, and DCs. SMAD4 loss resulted in an increase in the number and proportion of B220+ cells (40.5% vs 29.5%, *P =* .002; Supplementary Figure 5A). The absolute numbers of CD3+ T cells and CD11c+ DCs were significantly increased in GALTs of *Smad4*^*ΔLrig1*^*Ccr6*^*GFP/+*^mice compared to SMAD4+ *Ccr6*^*GFP/+*^ mice (Supplementary Figure 5B-C). However, the proportion of these cell types within GALTs were unchanged (data not shown). B cell, T cell, and DC distributions were not altered within the GALTs. B cells remained localized to the germinal center of GALTs, while T cells were spread throughout the GALT periphery, and DCs were localized to the subepithelial dome of GALTs, regardless of CCR6 expression status (Supplementary Figure 5A-C).

### Smad4 loss *increases immune cell infiltration in mouse colonic stroma, particularly T cell subsets and CD11c+ DCs, that is abrogated by additional loss of Ccr6*

To understand how epithelial *Smad4* loss impacts recruitment of specific immune cell types and to what degree these changes depend on functional CCL20/CCR6 signaling, we used flow cytometry to quantify relative changes in leukocyte populations, distinguishing epithelial-associated immune cells from those in the sub-epithelial stroma. Colitis was induced by administration of three rounds of DSS in six SMAD4+ *Ccr6*^*GFP/+*^, five SMAD4+ *Ccr6*^*GFP/GFP*^, five *Smad4*^*ΔLrig1*^ *Ccr6*^*GFP/+*^, and four *Smad4*^*ΔLrig1*^ *Ccr6*^*GFP/GFP*^ mice. Parallel myeloid and lymphoid flow cytometry staining panels were used to characterize the compositions of each compartment one month after the end of DSS treatment.

Within the colonic stroma, loss of *Ccr6* alone had little effect on the immune populations examined. However, a number of changes in immune cell composition were observed with epithelial cell loss of *Smad4. Smad4*^*ΔLrig1*^ *Ccr6*^*GFP/+*^ mice demonstrated a greater than two-fold increase in proportion of CD45+ immune cells in the colonic stromal tissues when compared to SMAD4+ *Ccr6*^*GFP/+*^ mice (41.4% vs 19.6%, *P <*.001) and this increase was abrogated by loss of *Ccr6* (*Smad4*^*ΔLri1*^ *Ccr6*^*GFP/+*^ mice 41.4% vs. *Smad4*^*ΔLrig1*^*Ccr6*^*GFP/GFP*^ mice 15.6%%, *P <*.001; Figure 3A-B). After normalizing to the CD45+ population, loss of *Smad4* alone resulted in increased CD3+ T cells. Among CD3+ T cells, the proportion of CD4+ T cells, CD4+ FoxP3+ Tregs, and CD4+ IL-17a+ cells were increased in the context of *Smad4* loss, but these increases were dependent upon *Ccr6* expression (Figure 3C-E). *Smad4*^*ΔLrig1*^*Ccr6*^*GFP/+*^ mice also had a significantly increased stromal CD11c+ population (Figure 3A, F). Further gating demonstrated that 60% of these cells were CD11b- and 52% were CX3CR1+, both of which were significantly increased in *Smad4*^*ΔLrig1*^ *Ccr6*^*GFP/+*^ mice as compared to SMAD4+ mice and these increases were also dependent upon *Ccr6* expression (Figure 3G). The expansion of these cell types in the absence of *Smad4* expression was likely due to recruitment of CCR6+ cells. Increases in CCR6+ cells marked by GFP+ staining within the CD45+, CD3+, CD4+, FoxP3+, IL-17a+, and CD11c+ populations mirrored the observed increases in these cell subtypes in *Smad4*^*ΔLrig1*^*Ccr6*^*GFP/+*^ mice (Supplementary Figure 6A). Additionally, the overwhelming majority of CCR6+ cells identified by GFP+ staining via flow cytometry in *Smad4*^*ΔLrig1*^*Ccr6*^*GFP/+*^ mice were immune cells (Supplementary Figure 6B).

**Figure 3.**
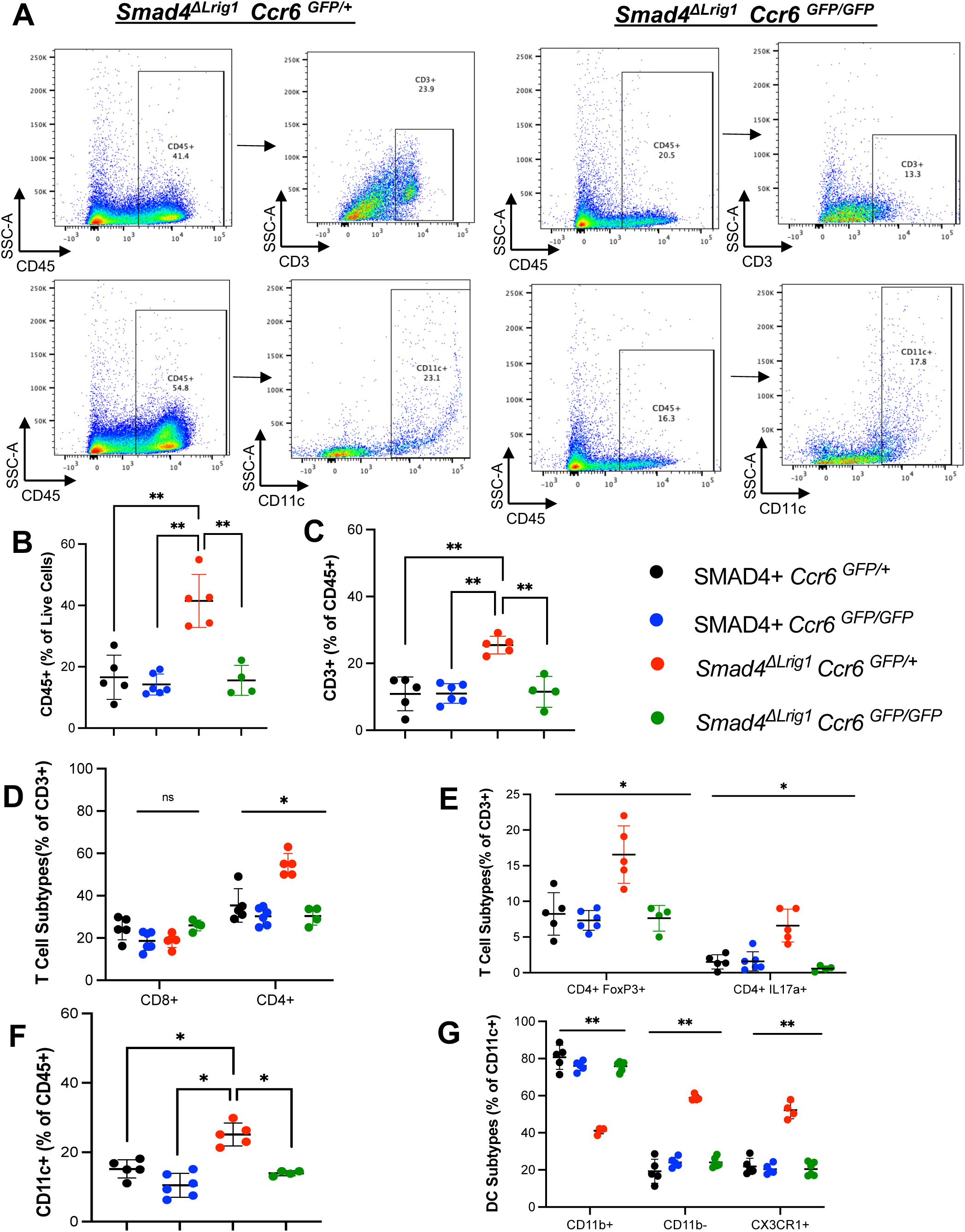
Loss of epithelial *Smad4* results in increased sub-epithelial immune cells in mouse colon in a CCR6-dependent manner. Colonic stroma was analyzed by flow cytometry 1 month after completion of DSS-induced colitis. **(A)** Representative flow cytometry scatter plots showing surface staining of colon stromal CD3+ or CD11c+ in CD45+ cells in a *Smad4*^*ΔLrig1*^ *Ccr6* ^*GFP/+*^ mouse and a *Smad4*^*ΔLrig1*^ *Ccr6* ^*GFP/GFP*^ mouse. Numbers in plot indicate percentage of cells in each gate. **(B)** Proportion of live stromal cells that were CD45+. **(C)** Proportion of CD45+ cells that were CD3+. **(D)** Proportion of CD3+ cells stained for CD8, CD4, **(E)** CD4 FoxP3, or CD4 IL17a. **(F)** Proportion of CD45+ cells that were CD11c+. **(G)** Proportion of CD11c+ cell stained for CD11b or CX3CR1 (\**P* < .05, ** *P* < .001).

To summarize the above findings, loss of epithelial SMAD4 expression leads to increased leukocyte recruitment to the colonic stroma, including CD4+ T cells, Tregs, T_h_17 cells, CD11c+ CD11b-DCs, and CD11c+ CX3CR1+ DCs (Supplementary Table 4). Notably, these changes are dependent on CCR6 expression, indicating that an intact CCL20/CCR6 axis is necessary to induce the stromal pro-inflammatory response observed due to epithelial SMAD4 loss.

### Ccr6 *mediates* Smad4-*dependent recruitment of mouse colon epithelial-associated T*_*h*_*17 and CD11c+ DCs*

Flow cytometric analysis was also performed on epithelial-associated cells. As in the sub-epithelial stroma, there was little effect on most of the examined populations in response to *Ccr6* loss alone, with one exception. *Ccr6* null mice demonstrated an expansion of colon epithelial-associated CD8+ T cells regardless of SMAD4 status, thus indicating a critical role for CCR6 in limiting CD8+ T cell trafficking to the intestinal epithelium (Figure 4D). On the other hand, loss of SMAD4 expression altered the proportion of specific sets of inflammatory cells associated with the epithelium. *Smad4*^*ΔLrig1*^ *Ccr6*^*GFP/+*^ mice demonstrated a significant increase in CD45+ cells within the colon epithelium fraction relative to SMAD4+ *Ccr6*^*GFP/+*^ mice (20.1% vs 9.6%, *P <* .001) and this increase was dependent upon *Ccr6* expression (*Smad4*^*ΔLrig1*^ *Ccr6*^*GFP/+*^ mice 20.1% vs. *Smad4*^*ΔLrig1*^ *Ccr6*^*GFP/GFP*^ mice 10.5%; *P <*.001; Figure 4A-B). There were no observed differences in total epithelial-associated CD3+ cells, CD4+ cells, or CD4+ FoxP3+ Tregs among the four genotypes, but the epithelium of *Smad4*^*ΔLrig1*^*Ccr6*^*GFP/+*^ mice exhibited significantly more CD4+ IL-17a+ cells than SMAD4+ *Ccr6*^*GFP/+*^ mice (6.4% vs 1.4%; *P <*.001) and loss of *Ccr6* abrogated this effect (*Smad4*^*ΔLri1*^ *Ccr6*^*GFP/+*^ mice 6.4% vs. *Smad4*^*ΔLrig1*^ *Ccr6*^*GFP/GFP*^ mice 1.2%; *P <*.001; Figure 4A,C-E). Thus, loss of epithelial SMAD4 expression in the colon results in a shift in CD3+ T cell composition towards increased T_h_17 cells.

**Figure 4.**
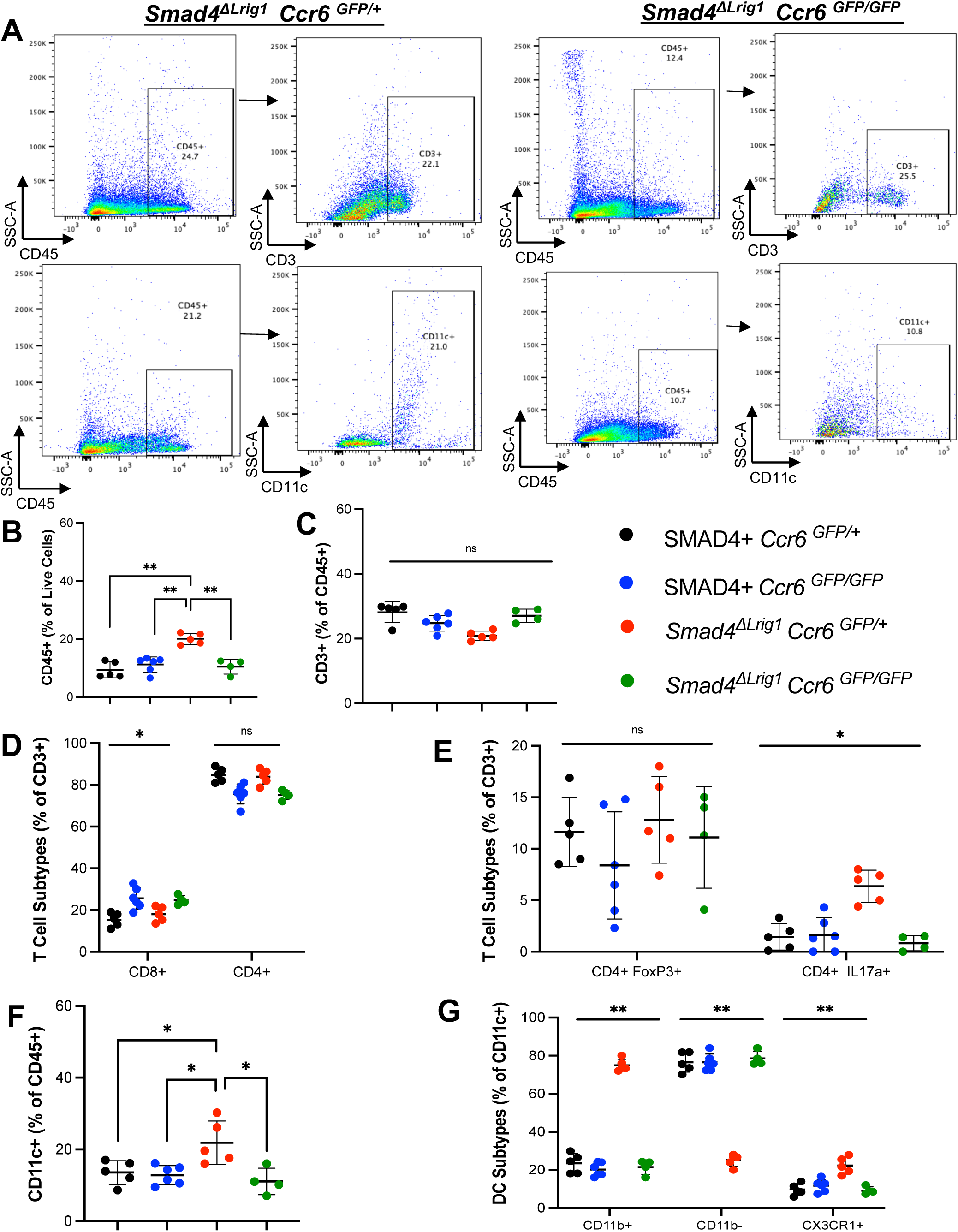
Loss of epithelial *Smad4* results in increased colon epithelial-associated immune cells in a CCR6-dependent manner. Cells isolated with colonic epithelium were analyzed by flow cytometry 1 month after completion of DSS-induced colitis. **(A)** Representative flow cytometry scatter plots showing surface staining for epithelial-associated CD3+ or CD11c+ in CD45+ cells of a *Smad4*^*ΔLrig1*^ *Ccr6* ^*GFP/+*^ mouse and a *Smad4*^*ΔLrig1*^ *Ccr6* ^*GFP/GFP*^ mouse. Numbers in plot indicate percentage of cells in each gate. **(B)** Proportion of live stromal cells that were CD45+. **(C)** Proportion of CD45+ cells that were CD3+. **(D)** Proportion of CD3+ cells stained for CD8, CD4, **(E)** CD4 FoxP3, or CD4 IL17a. **(F)** Proportion of CD45+ cells that were CD11c+. **(G)** Proportion of CD11c+ cell stained for CD11b or CX3CR1 (\**P* < .05, ** *P* < .001).

Among the myeloid cell populations, epithelial-associated CD11c+ DCs were increased in *Smad4*^*ΔLrig1*^ *Ccr6*^*GFP/+*^ mice relative to SMAD4+ *Ccr6*^*GFP/+*^ (21.9% vs 13.6%, *P =* .03), including a significant increase in the CD11c+ CX3CR1+ DC subset and these increases were also abrogated by loss of *Ccr6* (Figure 4A, F-G). The DC population observed within the epithelium differed from the stroma in that the majority of DCs associated with the epithelium were CD11c+ CD11b+, whereas the sub-epithelial stroma contained primarily CD11c+ CD11b-DCs. Further gating for GFP expression among these cell types demonstrated a concomitant increase in GFP+ cells among the increased T_h_17 and dendritic cell populations (Supplementary Figure 7A). Similar to stroma, the increase in CCR6+ cells seen with colon epithelial cell SMAD4 loss were overwhelmingly CD45+ (Supplementary Figure 7B).

In summary, loss of colon epithelial cell SMAD4 leads to increased recruitment of specific immune cell subsets within the epithelium, particularly T_h_17 cells and DCs including CD11c+ CD11b+ and CD11c+ CX3CR1+ DC subtypes that is dependent on CCR6 activity (Supplementary Table 4). *Ccr6* loss led to an increase in CD8+ IELs regardless of SMAD4 expression status as previously published ^13, 23^.

### CAC susceptibility due to SMAD4 loss is dependent on the CCL20/CCR6 axis

We previously published that loss of epithelial SMAD4 in the colon significantly predisposes mice to CAC within three months of DSS-induced chronic inflammation, and that these tumors are morphologically similar to human UCACs ^5^. To determine if this cancer susceptibility phenotype also depends on an intact CCL20/CCR6 axis, we induced chronic colitis in SMAD4+ *Ccr6*^*GFP/+*^, SMAD4+ *Ccr6*^*GFP/GFP*^, *Smad4*^*ΔLrig1*^ *Ccr6*^*GFP/+*^, and *Smad4*^*ΔLrig1*^ *Ccr6*^*GFP/GFP*^ mice with 3 cycles of DSS. Three months after completion of DSS treatment, 100% (8 of 8) of *Smad4*^*ΔLrig1*^ *Ccr6*^*GFP/+*^ mice developed adenocarcinomas of the colon with invasion deep into the submucosa, consistent with prior observations from our group ^5^. In contrast, only 18.2% (2 of 11) *Smad4*^*ΔLrig1*^ *Ccr6*^*GFP/GFP*^ mice developed these invasive cancers, one of which was characterized as a single invasive crypt (Figure 5A, C). Thus, loss of CCR6 expression is associated with a significant decrease in susceptibility to CAC development in *Smad4* null mice (100.0 vs 18.2%, *P <* .001) Loss of *Ccr6* expression was similarly associated with fewer invasive tumors per colon. The range of invasive adenocarcinomas for *Smad4*^*ΔLrig1*^ *Ccr6*^*GFP/+*^ mice was 1 – 3 tumors compared to 0 – 2 invasive adenocarcinomas per colon in *Smad4*^*ΔLrig1*^*Ccr6*^*GFP/GFP*^ mice (Figure 5B). We did note a single invasive crypt on a single section of one SMAD4+ *Ccr6*^*GFP/GFP*^ mouse, consistent with previous observations that DSS-induced colitis can occasionally induce tumorigenesis in mice without loss of SMAD4 but at longer time points ^24^.

**Figure 5:**
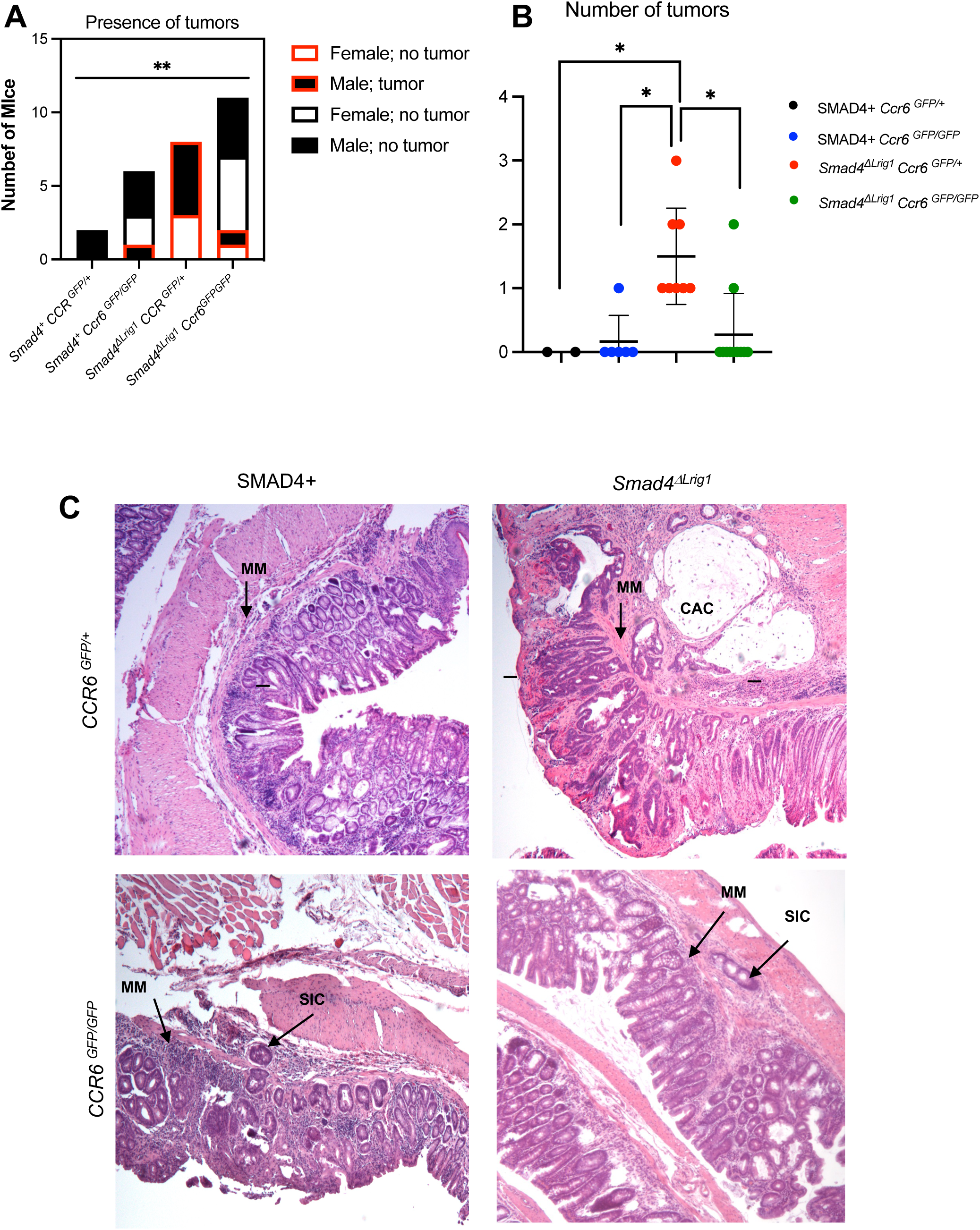
Loss of *Ccr6* prevents tumorigenesis after DSS-induced colitis. Mice of the indicated genotypes were analyzed for invasive tumors 3 months after the last cycle of DSS. **(A)** Number of mice that developed CAC s in each group. **(B)** Number of invasive tumors noted per mouse in each genotype. **(C)** Representative images of invasive adenocarcinoma of the distal colon of a *Smad4*^*ΔLrig1*^ *Ccr6*^*GFP/+*^ mouse as well as the single invasive crypt seen in a *Smad4*^*ΔLrig1*^ *Ccr6*^*GFP/EGFP*^ and a *Smad4+ Ccr6*^*GFP/GFP*^ mouse (MM: muscularis mucosa; SIC: single invasive crypt; CAC: colitis associated cancer; Scale bars: 100μm; \**P* < .05, ** *P* < .001).

Tissue sections from the same mice were subjected to histologic grading of colitis. Among the *Smad4*^*ΔLrig1*^ mice, there was no significant difference between *Ccr6* expressing and *Ccr6* null mice in terms of weight loss during DSS or overall inflammation score at the time of tumor scoring (data not shown). However, in SMAD4+ control mice, *Ccr6* expression was associated with a higher percentage of crypts involved with colitis (2 vs 1.17, *P =* .03) and deeper crypt damage (2 vs 1.67, *P =* .004) compared to *Ccr6* null mice. Loss of *Smad4* also decreased the severity of colitis; SMAD4+ *Ccr6*^*GFP/+*^ mice had higher average crypt damage scores than *Smad4*^*ΔLrig1*^ *Ccr6*^*GFP/+*^ mice (2 vs 0.5, *P <*.001) and a similar relationship was observed between SMAD4+ *Ccr6*^*GFP/GFP*^ and *Smad4*^*ΔLrig1*^ *Ccr6*^*GFP/GFP*^ mice (1.67 vs 0.46, *P =* .002; Supplementary Table 5).

Thus, loss of *Smad4* reduced histologic evidence of the severity of ongoing colitis yet contributed to the development of invasive tumors at a higher rate and frequency. These findings indicate that the development of CAC in *Smad4*^*ΔLrig1*^ *Ccr6*^*GFP/+*^mice is not solely dependent on the severity of colitis alone, but rather on the unique response to colitis seen with epithelial SMAD4 loss, perhaps related to specific leukocyte recruitment.

### Altered immune cell landscape in the setting of SMAD4 loss persists with tumorigenesis

The above data suggest that loss of epithelial SMAD4 expression promotes a unique immune microenvironment within one month of induction of chronic colitis and leads to a profound susceptibility to CAC development by three months, both of which are dependent on CCR6 expression. To determine if the observed immune cell changes persist at the time of tumorigenesis, we performed IHC of specific immune cell subtypes in SMAD4+ *Ccr6*^*GFP/+*^, SMAD4+ *Ccr6*^*GFP/GFP*^, *Smad4*^*ΔLrig1*^ *Ccr6*^*GFP/+*^, and *Smad4*^*ΔLrig1*^ *Ccr6*^*GFP/GFP*^ mice 3 months after completion of DSS treatment. Overall, the unique immunophenotypic changes observed in *Smad4*^*ΔLrig1*^ *Ccr6*^*GFP/+*^ mice one month after DSS completion persisted at three months in regions adjacent to tumors. However, the spatial distributions of immune cell subtypes 3 months after induction of colitis were largely unchanged with or without loss of *Smad4* or *Ccr6*. CD3+ cells were increased throughout the GALTs with loss of *Smad4* only when *Ccr6* was expressed. Within the epithelium, a similar number of CD3+ T cells were observed with or without loss of *Smad4* or *Ccr6* (Supplementary Figure 8A). Examination of CD8+ IELs by IHC at three months corroborated our flow cytometry results that *Ccr6* loss leads to increased CD8+ IELs regardless of SMAD4 status (Supplementary Figure 8B).

FoxP3+ Tregs were noted to be localized to GALTs or within the epithelium, without significant distribution through the rest of the stroma (Figure 6A). Within the GALTs, FoxP3+ Tregs were markedly increased in *Smad4*^*ΔLrig1*^ *Ccr6*^*GFP/+*^ mice but not in *Smad4*^*ΔLrig1*^ *Ccr6*^*GFP/GFP*^ mice (Figure 6A). Similar to mice at one month after DSS treatment, there was no observable difference in epithelial-associated Tregs (Figure 6B). IL17+ cells were notably increased in the GALTs and epithelium following *Smad4*^*ΔLrig1*^ loss but only in the presence of *Ccr6* expression. (Figure 7A-B.) The distribution of CD11c+ DCs was similar among mice from all four genotypes (Supplementary Figure 9). Together, our findings indicate that epithelial SMAD4 loss induces immunophenotypic changes that persist from 1 month after the DSS treatment, when no tumors are detectable, through 3 months after DSS when invasive cancers are prevalent in all of the *Smad4*^*ΔLrig1*^ *Ccr6*^*GFP/+*^mice. Both the immune cell changes and tumorigenesis are blocked by loss of *Ccr6*, linking the immune phenotype to tumorigenesis.

**Figure 6.**
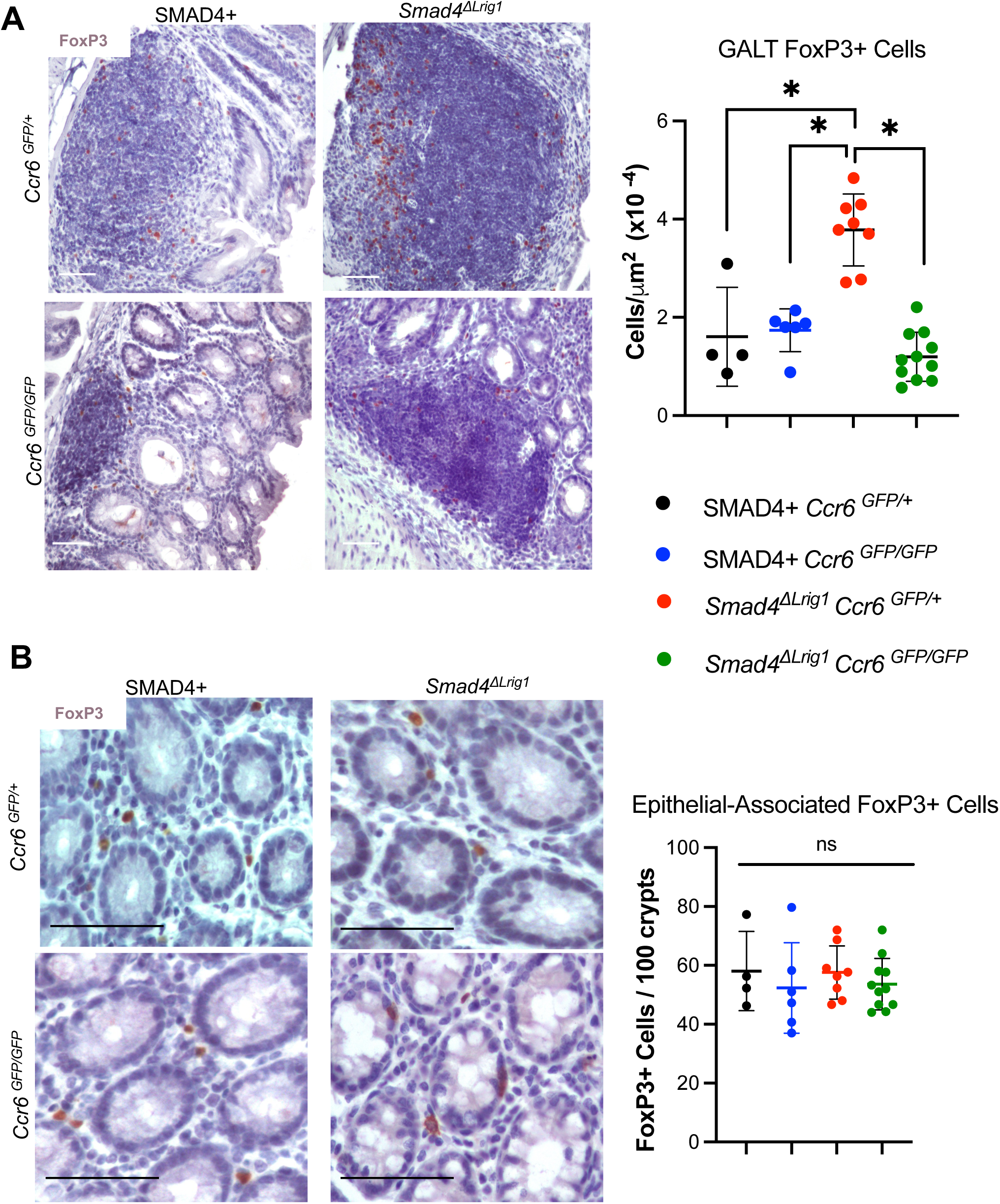
Loss of epithelial *Smad4* resulted in significantly increased FoxP3+ cells in GALTs only in the presence of CCR6. Staining for FoxP3 (brown) in representative samples for **(A)** GALTs or **(B)** crypt-associated regions comparing mice of the indicated genotypes (scale bars: 100μm). Graphs represent quantification of staining. Data points represent mean quantification per mouse (\**P* < .05).

**Figure 7.**
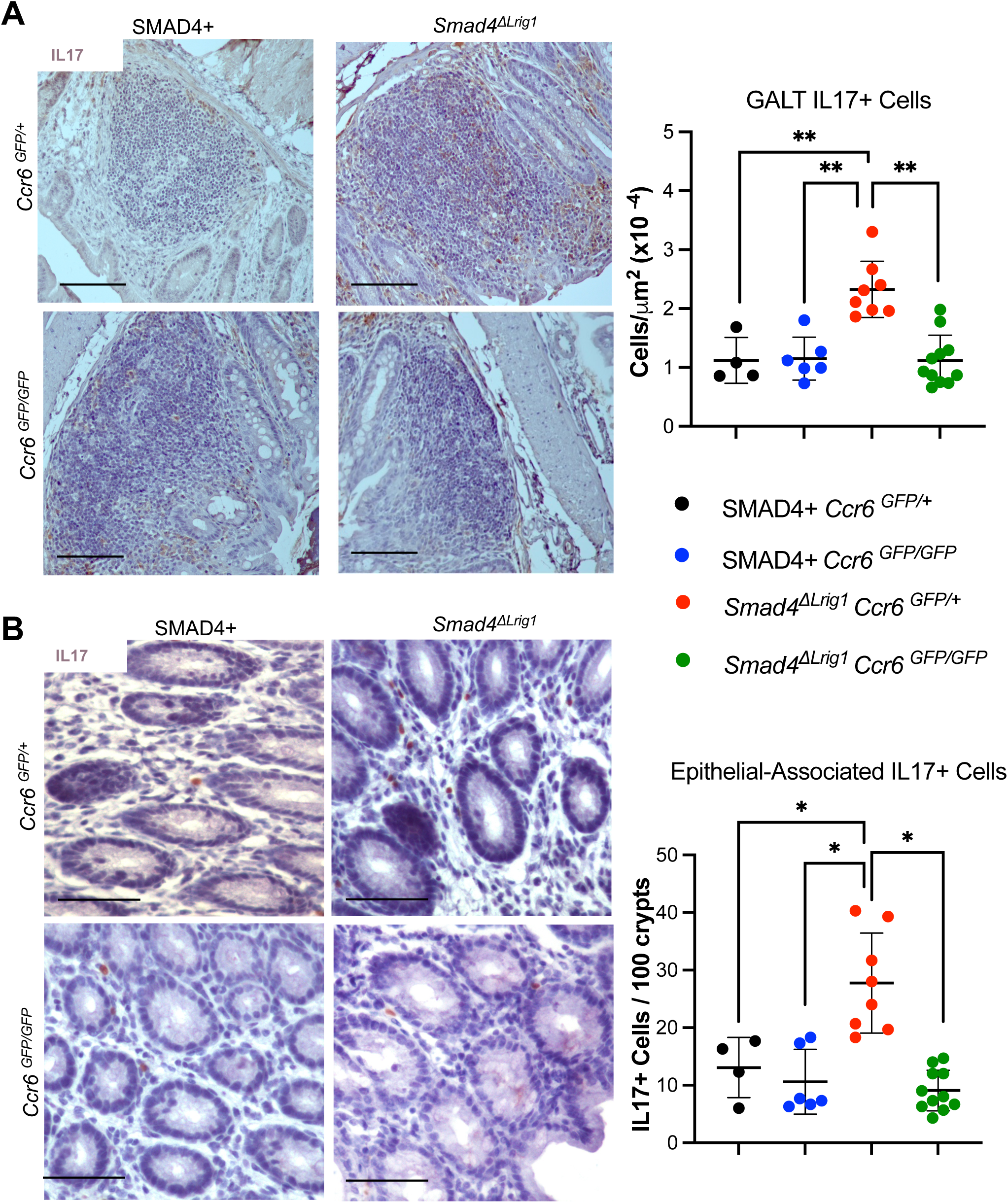
*Smad4* loss resulted in increased IL17+ cells in GALTs and crypt-associated regions only in the presence of CCR6. Staining for IL17+ cells (brown) **(A)** within GALTs and **(B)** juxtaposed to epithelium comparing colons of mice of the indicated genotypes (scale bars: 100μm). Graphs represent quantification of staining. Data points represent mean quantification per mouse (\**P* < .05, ** *P* < .001).

## Discussion

Multiple defects in the TGFβ signaling pathway, including receptor mutations and mutations in *SMAD genes*, most frequently *SMAD4*, have been previously implicated in multiple gastrointestinal pathologies including human IBD and CRC ^4, 25, 26^. Previous attention has primarily focused on the roles of TGFβ in maintenance of epithelial homeostasis including regulation of cell growth, cell cycle, differentiation, and apoptosis, all of which likely factor into the tumor suppressor effects of TGFβ family signaling ^27, 28^. Our data support a novel tumor suppressor function of the canonical TGFβ signaling pathway via modulation of inflammatory responses in the epithelium. It is already well appreciated that the TGFβ signaling pathway plays an important immunosuppressive role through its direct actions on immune cells, including inhibition of T_h_1 helper cell, cytotoxic T cell, and NK cell responses, and simultaneous promotion of Treg differentiation and function.^29, 30^ Here, we add another level of homeostatic immune regulation by TGFβ signaling via direct effects on epithelium, downregulating epithelial inflammatory genes, particularly *Ccl20*, to suppress specific mucosal inflammatory responses and inflammation-associated carcinogenesis.

Several studies have implicated defective epithelial TGFβ signaling or the CCL20/CCR6 axis in IBD and CRC development ^31, 32^. However, we have identified for the first time a significant inverse correlation between *SMAD4* and *CCL20* expression in human IBD, colon cancer, and rectal cancer specimens. By utilizing mouse models, we have been able to link the epithelial function of SMAD4 to regulation of CCL20/CCR6 signaling, suggesting a mechanism underlying inflammation and tumorigenesis downstream of decreased or lost TGFβ/SMAD4 signaling in human patients along the IBD to CAC spectrum.

Concurrent with increased chemokine/cytokine expression in *Smad4* null mouse colon epithelium is an increase in pro-inflammatory signaling within the underlying stroma. Many of the upregulated genes in the stroma have known roles in the innate and adaptive immune responses, including those involved with mucosal immune trafficking and GALT structure and function. Additionally, we observed evidence of increased GALT size in *Smad4*^*ΔLrig1*^ mice. CCL20 signaling to CCR6 is known to regulate immune cell trafficking to mucosa-associated lymphoid structures such as GALTs ^33^. Consistent with this concept, not only did GALT size increase with increased CCL20 expression after loss of epithelial SMAD4 loss, but this increase in GALT size was abrogated in mice without *Ccr6* expression. Increased *Ccl20* expression, *Ccr6* expression, and GALT size, and upregulation of genes that function in pathogen recognition (*Tlr9, Pyhin2, Fcrla, Ly86*) support a role for the SMAD4-mediated regulation of CCL20 in modulating mucosal immune responses through GALT activity.

Herein we demonstrated that much of the inflammatory response to *Smad4* loss in the epithelium can be abrogated by loss of *Ccr6*, which is normally expressed on many of the immune cells that are increased with epithelial SMAD4 loss. The reduction in tumor burden in response to *Ccr6* deletion in *Smad4*^*ΔLrig1*^ mice suggests the CCL20/CCR6 axis is one of the most critical inflammatory pathways downstream of TGFβ signaling in the development of CAC. Our findings are consistent with previous findings that loss of *Ccr6* expression protects against *APC* mutant-driven and azoxymethane (AOM)/DSS-driven tumorigenesis in mice ^34^. While restoring expression of tumor suppressor genes remains clinically intractable, targeting downstream pathways such as CCL20/CCR6 may be one approach for more selective clinically targeted therapies.

Our findings additionally provide insight into immune cell subtypes downstream of SMAD4 loss that could be targeted for improved therapies for prevention of or treatment of CACs. Our results suggest that targeted therapy against CCR6 may be effective in preventing CAC in ulcerative colitis patients. There may also be efficacy in targeting specific cell types that respond to CCR6 activation. In particular, we observed an increase in stromal Tregs, T_h_17 cells, and DCs as well as epithelial-associated T_h_17 cells and DCs in *Smad4*^*ΔLrig1*^ mice (summarized in Supplementary Figure 7). Tregs are involved in suppressing immune responses, particularly against commensal bacteria in the gut. However, the immunomodulatory role of Tregs may hinder tumor immunosurveillance and thereby contribute to tumor formation ^35^. Several studies have shown that Tregs are more abundant in inflamed IBD colon mucosa and in colon tumors compared to non-inflamed or healthy colon samples in humans ^36, 37^. We observed increased Tregs in the stroma of *Smad4*^*ΔLrig1*^ mice after induction of inflammation at the time of CAC formation and before tumors developed, suggesting a potential role of Tregs in CAC tumorigenesis.

In the present study, we also observed an increased number of T_h_17cells in both the stromal and epithelial compartments of *Smad4*^*ΔLrig1*^ mice. While the role of T_h_17 cells in tumor immunity remains poorly understood, immunoneutralization of IL-17 has been shown to inhibit colitis and CAC formation in an enterotoxigenic colitis mouse model ^38^. Thus, two CCR6+ immune cells, T_h_17 cells and FoxP3+ Tregs, are candidate effector cell types which may play a causative role in colitis-associated tumorigenesis due to SMAD4 loss and CCL20 secretion. Additionally, we also observed increased CD11c+ CX3CR1+ cells in the epithelial and stromal compartment of *Smad4*^*ΔLrig1*^ mice with intact CCR6 expression. CD11c+ CX3CR1+ cells have previously been observed in the mucosa of IBD patients and are known to produce both IL-1 and IFNβ, which are required for T_h_17 cell differentiation and Treg expansion, respectively ^39, 40^. Future studies will define which of these CCR6+ cell types are critical to tumorigenesis and whether they could be targeted clinically in the setting of colitis.

In summary, we have identified a specific chemokine pathway, epithelial CCL20 signaling to CCR6 on immune cells, that is normally inhibited by TGFβ family canonical signaling under homeostatic conditions but is activated by epithelial loss of SMAD4 in the colon and is essential for the development of CAC in conditional *Smad4* knockout mice. We have also identified the specific inflammatory cells that respond to that chemokine axis and correlate with carcinogenesis. Future research will identify how specific immune cell functional alterations in the context of epithelial loss of SMAD4 expression contribute to colon epithelial cell transformation and cancer progression and whether these represent potential opportunities in the prevention or treatment of colorectal cancer.

## Abbreviations

CRC: colorectal cancer
IBD: inflammatory bowel disease
TGFβ: transforming growth factor β
CAC: colitis-associated carcinoma
CCL20: c-c motif chemokine ligand 20
CCR6: c-c motif chemokine receptor 6
GALT: gut-associated lymphoid tissue
DSS: dextran sodium sulfate
TMA: tissue microarray
UCAC: ulcerative colitis-associated cancer
IHC: immunohistochemistry
IF: immunofluorescent
UCa: active ulcerative colitis
CDa: active Crohn’s disease
HC: healthy colon
UCi: inactive ulcerative colitis
TCGA: the cancer genome atlas
CC: colon cancer
HCc: healthy colon control
RC: rectal cancer
MxIF: multiplex immunofluorescence
FAE: follicle-associated epithelium

**Author names in bold designate shared co-first authorship**

## Supplementary Figure Legends

**Supplementary Figure 1.**
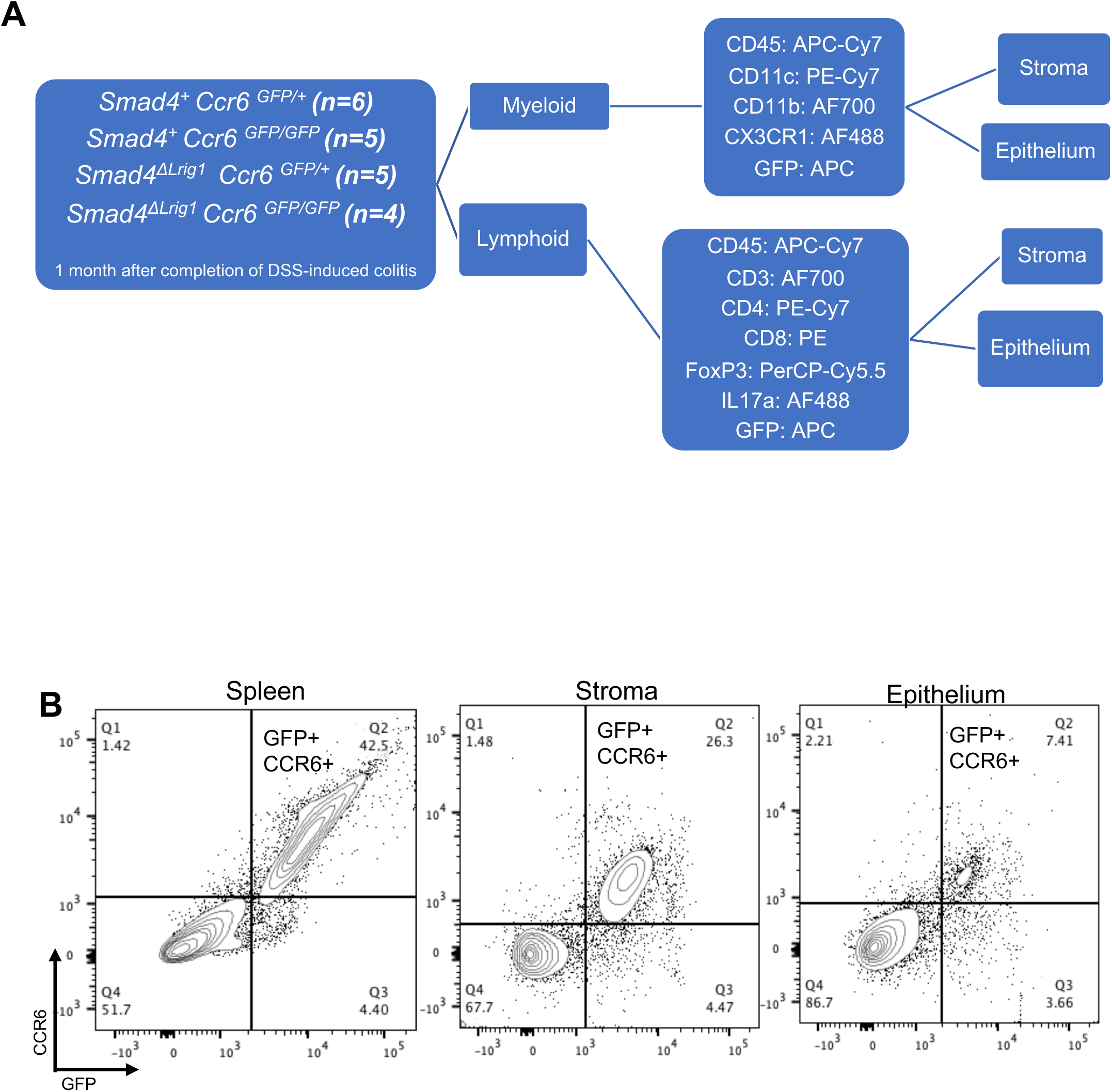
Schematic diagram of flow cytometry experiment design. **(A)** Mice were sacrificed 1 month after completion of DSS treatment. Cells were isolated from the epithelial and stromal compartments from each mouse, stained with separate myeloid and lymphoid antibody panels, and analyzed by flow cytometry. **(B)** Cells were stained and analyzed by flow cytometry for CCR6 and GFP to assess the accuracy of using GFP as a surrogate for *Ccr6* promoter activity in mouse spleen, colonic stroma, and colonic epithelial region. Numbers indicate percentage of cells in each gate.

**Supplementary Figure 2.**
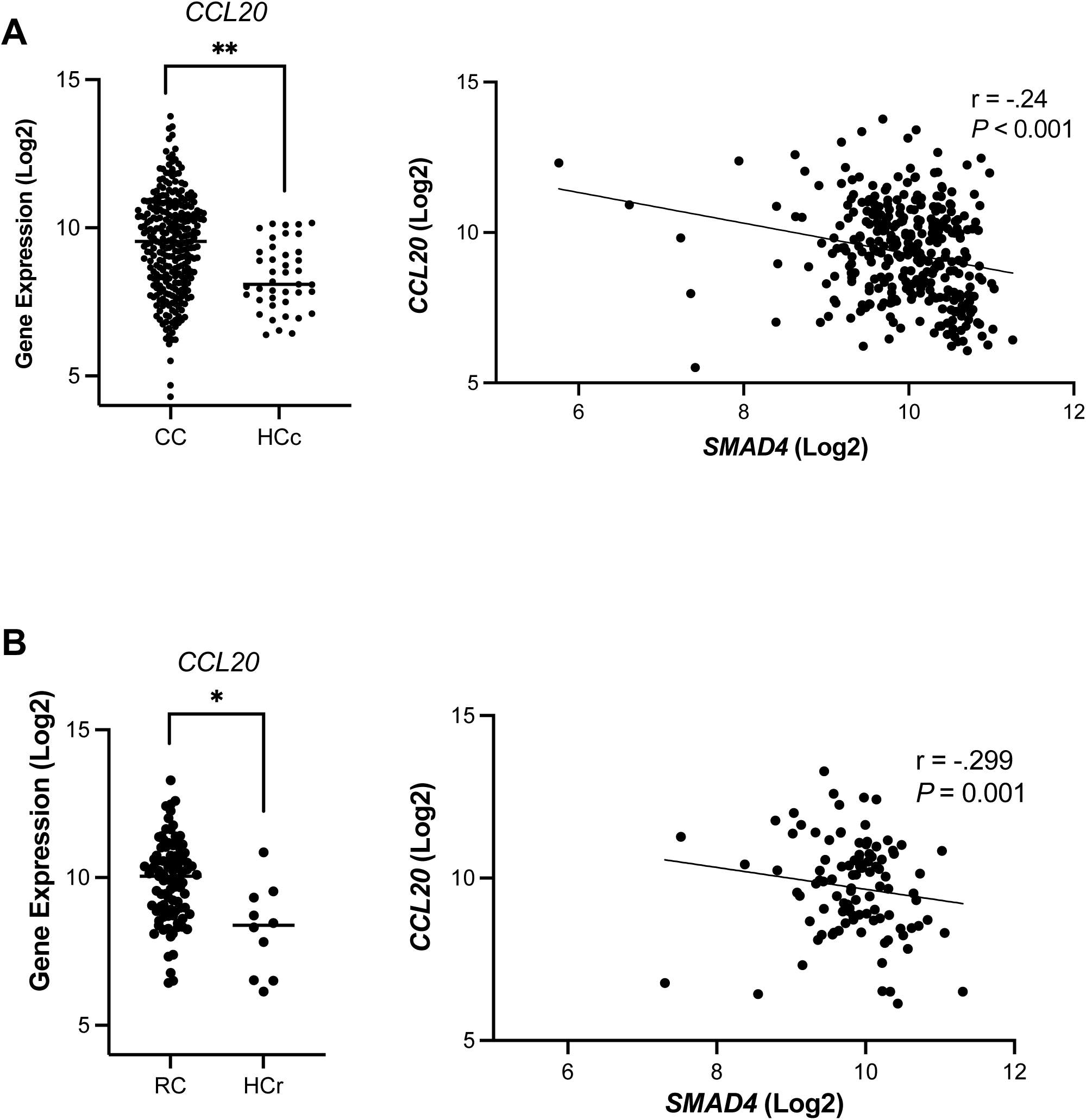
*SMAD4* expression is negatively correlated with *CCL20* expression in human sporadic colon and rectal cancer specimens. **(A-B)** *In silico* analysis of the TCGA database for **(A)** colon and **(B)** rectal cancer (CC: colon cancer; HCc: healthy colon control; RC: rectal cancer; HCr: healthy rectum control; \**P <* .05, \*\**P <* .001).

**Supplementary Figure 3.**
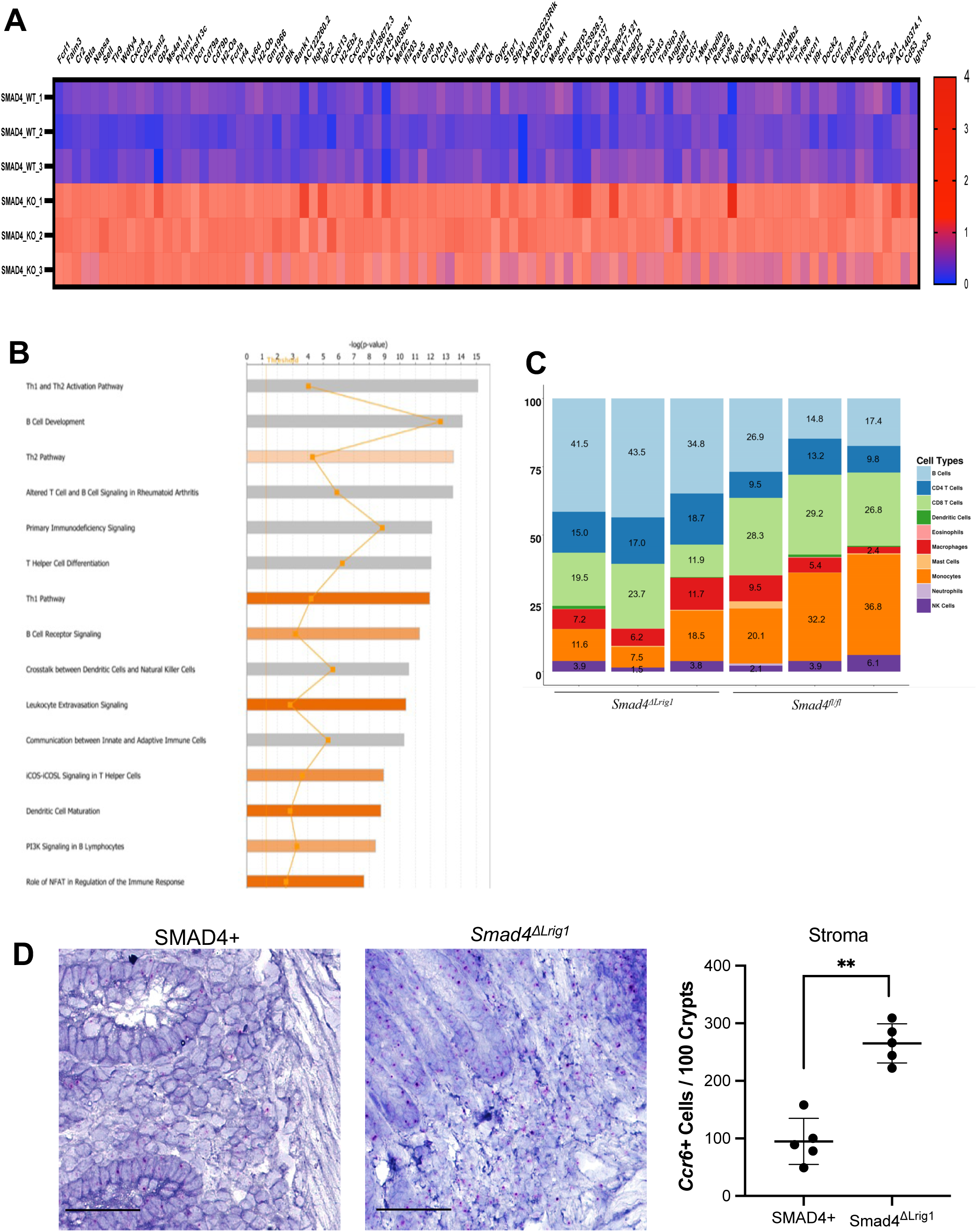
Inflammation-associated genes, including *Ccr6*, are upregulated in the colonic stroma of *Smad4*^*ΔLrig1*^ mice. **(A)** Heat map showing differential expression of 95 genes, including *Ccr6*. **(B)** Ingenuity pathway analysis was used to identify pathways significantly altered in the surrounding stroma by loss of epithelial SMAD4. **(C)** ImmuCC computational analysis was used to infer immune cell composition in *Smad4*^*ΔLrig1*^ and SMAD4+ mice. **(D)** Representative RNAscope ® *in situ* hybridization of *Smad4*^*ΔLrig1*^ and SMAD4+ colons for *Ccr6* transcripts and quantification of cells expressing *Ccr6*. Data points represent mean quantification per mouse (\*\**P <* .001; scale bars = 100μm).

**Supplementary Figure 4.**
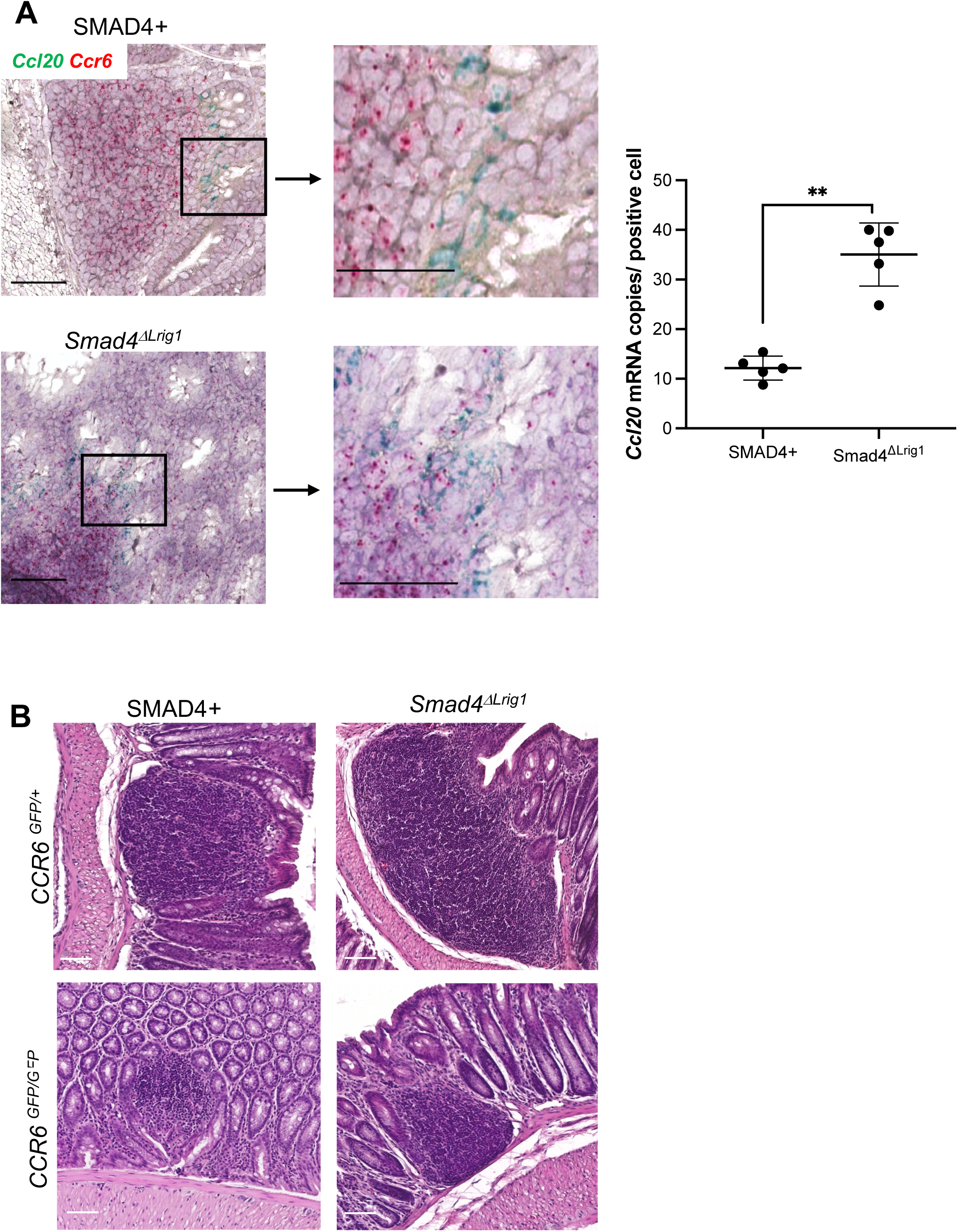
Loss of epithelial *Smad4* results in increased *Ccl20* expression in follicle-associated epithelial cells and increased GALT size. **(A)** Representative RNAscope ® *in situ* hybridization images for *Ccl20* and *Ccr6* transcripts and quantification of *Ccl20* staining in epithelial cells overlying GALTs in *Smad4*^*ΔLrig1*^ and SMAD4+ colons (\*\**P <* .001; *Ccl20 =* green, *Ccr6 =* red; scale bars = 100μm). **(B)** Representative H&E images of different GALT sizes based on *Smad4* and *Ccr6* expression as indicated. Data points represent mean quantification per mouse (scale bars: 100μm).

**Supplementary Figure 5.**
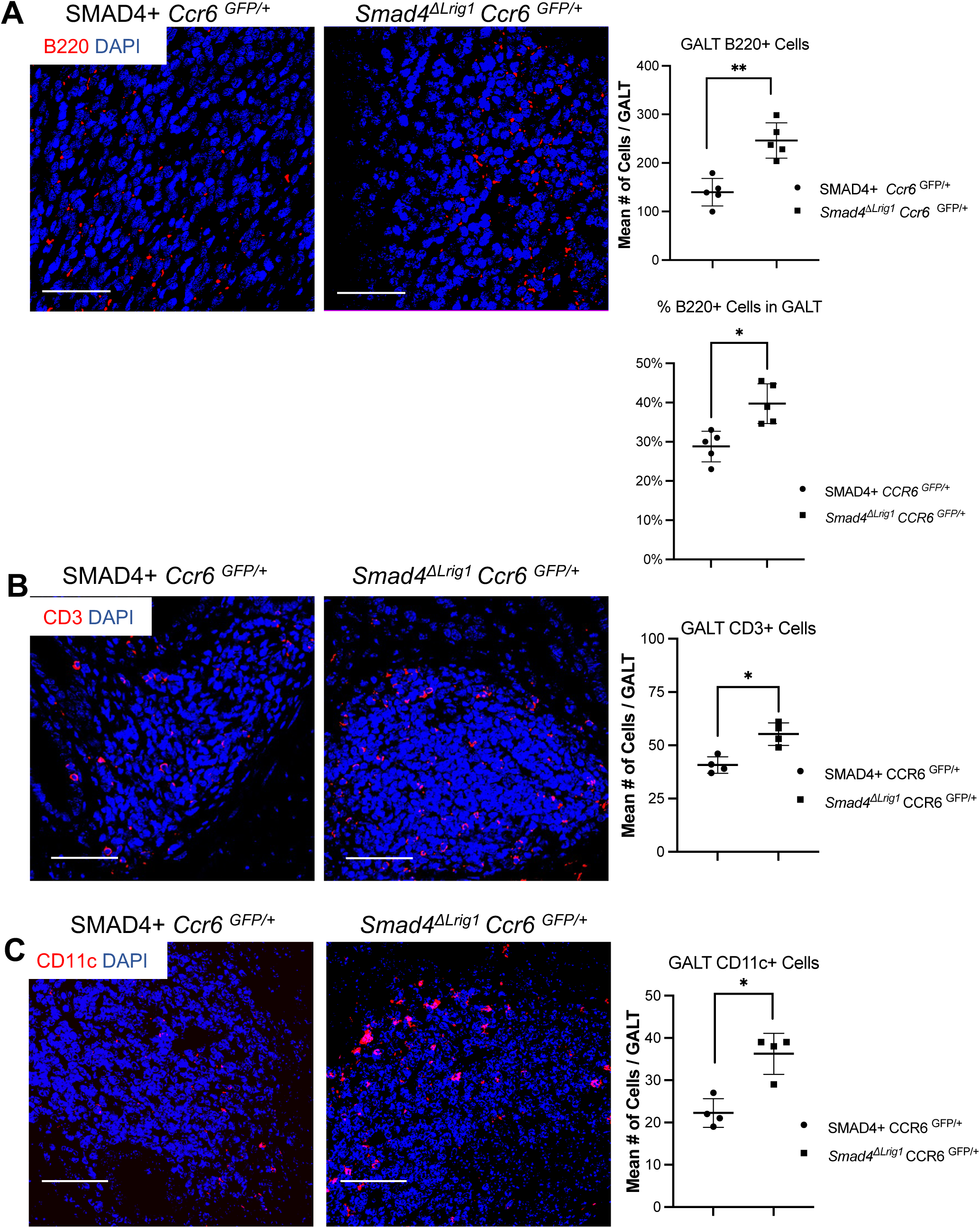
Loss of epithelial *Smad4* results in increased number of (A) B220+ B cells, (B) CD3+ T cells, and (C) CD11c+ dendritic cells under homeostatic conditions, but does not alter their spatial distributions. Representative IF images and quantification of cell types in GALTs of five *Smad4*^*ΔLrig1*^ and five SMAD4+ mice. Data points represent mean quantification per mouse (DAPI = blue; B220, CD3, CD11c = red; scale bars: 100*μm*).

**Supplementary Figure 6:**
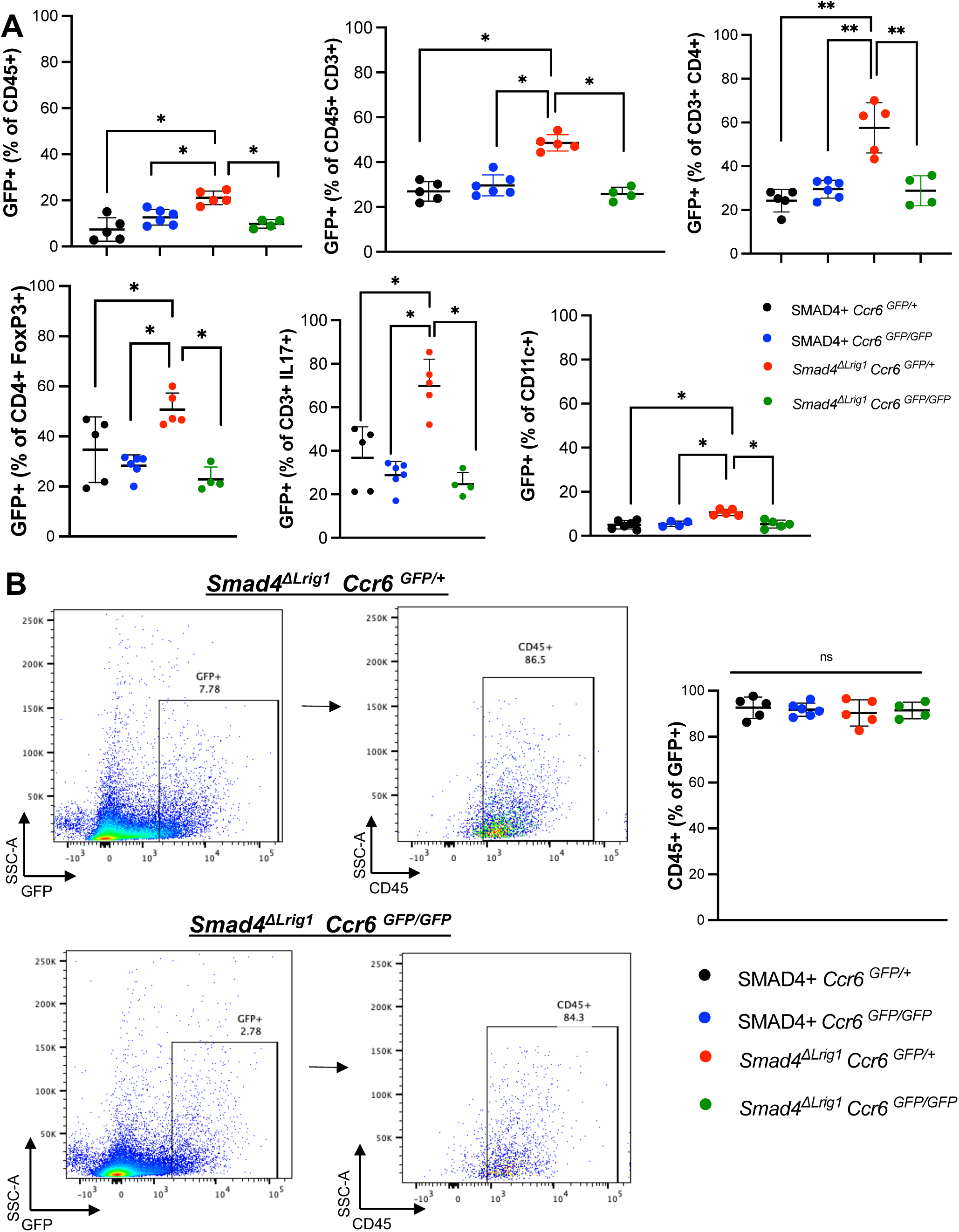
Flow cytometry analysis of *Smad4*^*ΔLrig1*^ *Ccr6*^*GFP/+*^ mice demonstrated that the observed increases in stromal immune cells were due to increased CCR6+ cell numbers. **(A)** Percentage of stromal CD45+, CD3+, CD4+, CD4+ FoxP3+, CD4+ IL17a+, and CD11c+ cells that were also GFP+. **(B)** Representative flow cytometry scatter plots and quantification showing the proportion of GFP+ cells that are CD45+ in the stroma of a *Smad4*^*ΔLrig1*^ *Ccr6* ^*GFP/+*^ mouse and a *Smad4*^*ΔLrig1*^ *Ccr6* ^*GFP/GFP*^ mouse. Numbers in plots indicate percentage of cells in each gate (*P <* .05, \*\**P <* .001).

**Supplementary Figure 7:**
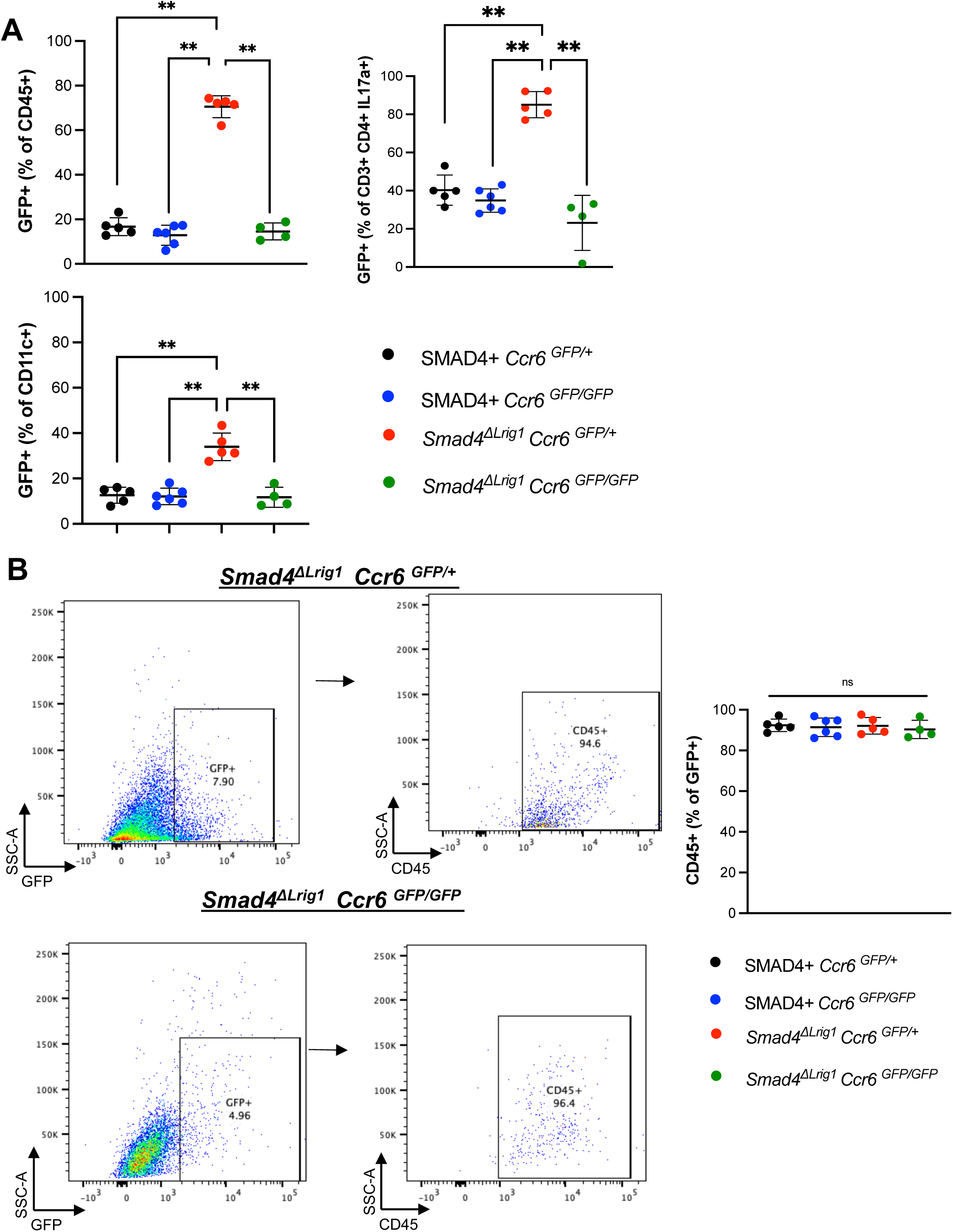
Flow cytometry analysis of *Smad4*^*ΔLrig1*^ *Ccr6*^*GFP/+*^ mice demonstrated that the observed increases in epithelial-associated immune cells were due to increased CCR6+ cell numbers. **(A)** Percentage of epithelial-associated CD45+, CD4+ IL17a+, and CD11c+ cells that were also GFP+. **(B)** Representative flow cytometry scatter plots and quantification showing the proportion of epithelial-associated GFP+ cells that are CD45+ in a *Smad4*^*ΔLrig1*^ *Ccr6* ^*GFP/+*^ mouse and a *Smad4*^*ΔLrig1*^ *Ccr6* ^*GFP/GFP*^ mouse. Numbers in plots indicate percentage of cells in each gate (*P <* .05, \*\**P <* .001).

**Supplementary Figure 8.**
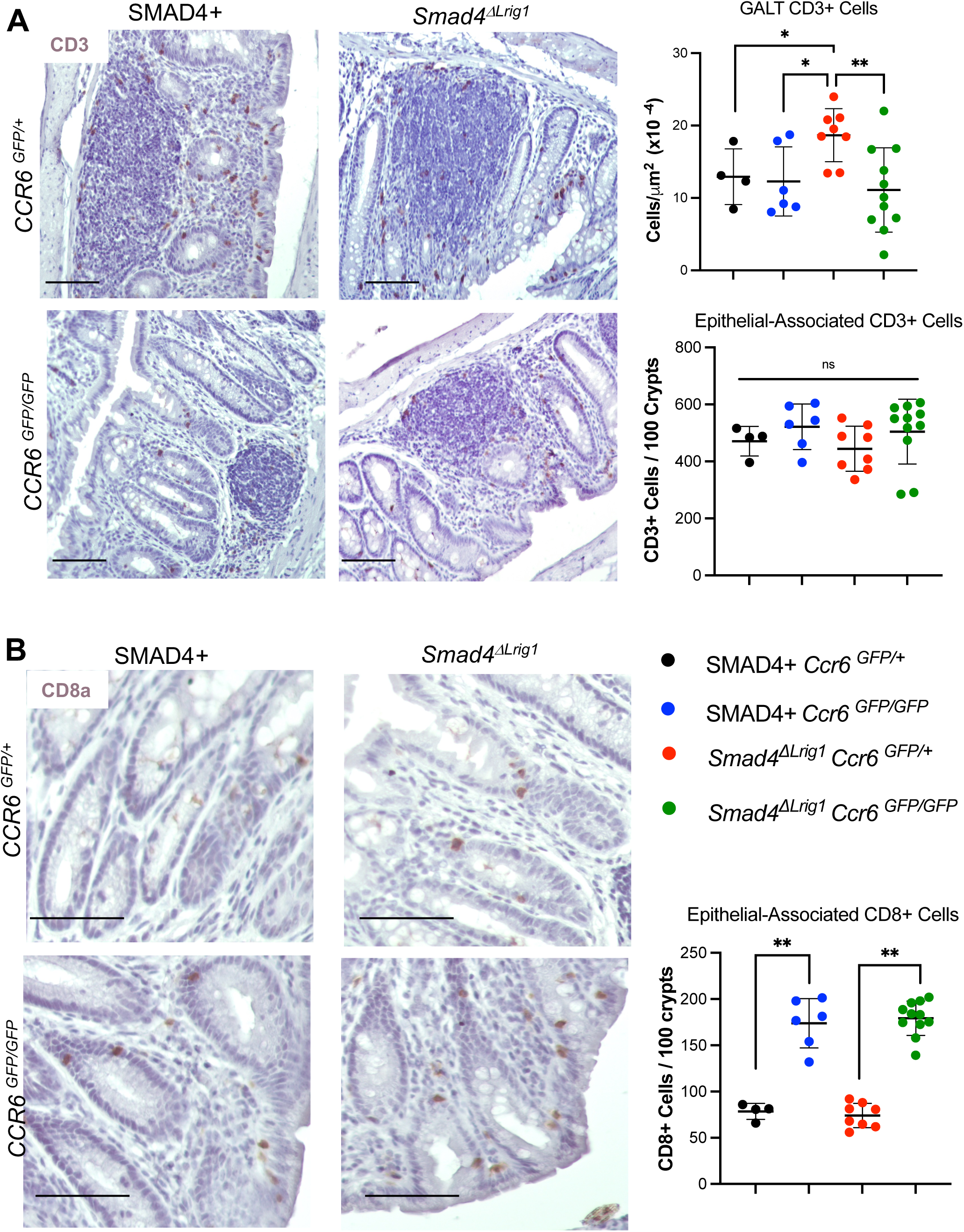
*Smad4* loss was associated with increased GALT CD3+ cells and loss of *Ccr6* led to an increase in CD8+ intraepithelial lymphocytes three months post-DSS exposure. **(A)** Representative images and quantification of CD3+ cells in GALTS in colons of indicted genotypes. **(B)** Representative images and quantification of CD8+ cells in crypt regions in colons of indicted genotype. Data points represent mean quantification per mouse (scale bars: 100μm; *P <* .05, \*\**P <* .001).

**Supplementary Figure 9.**
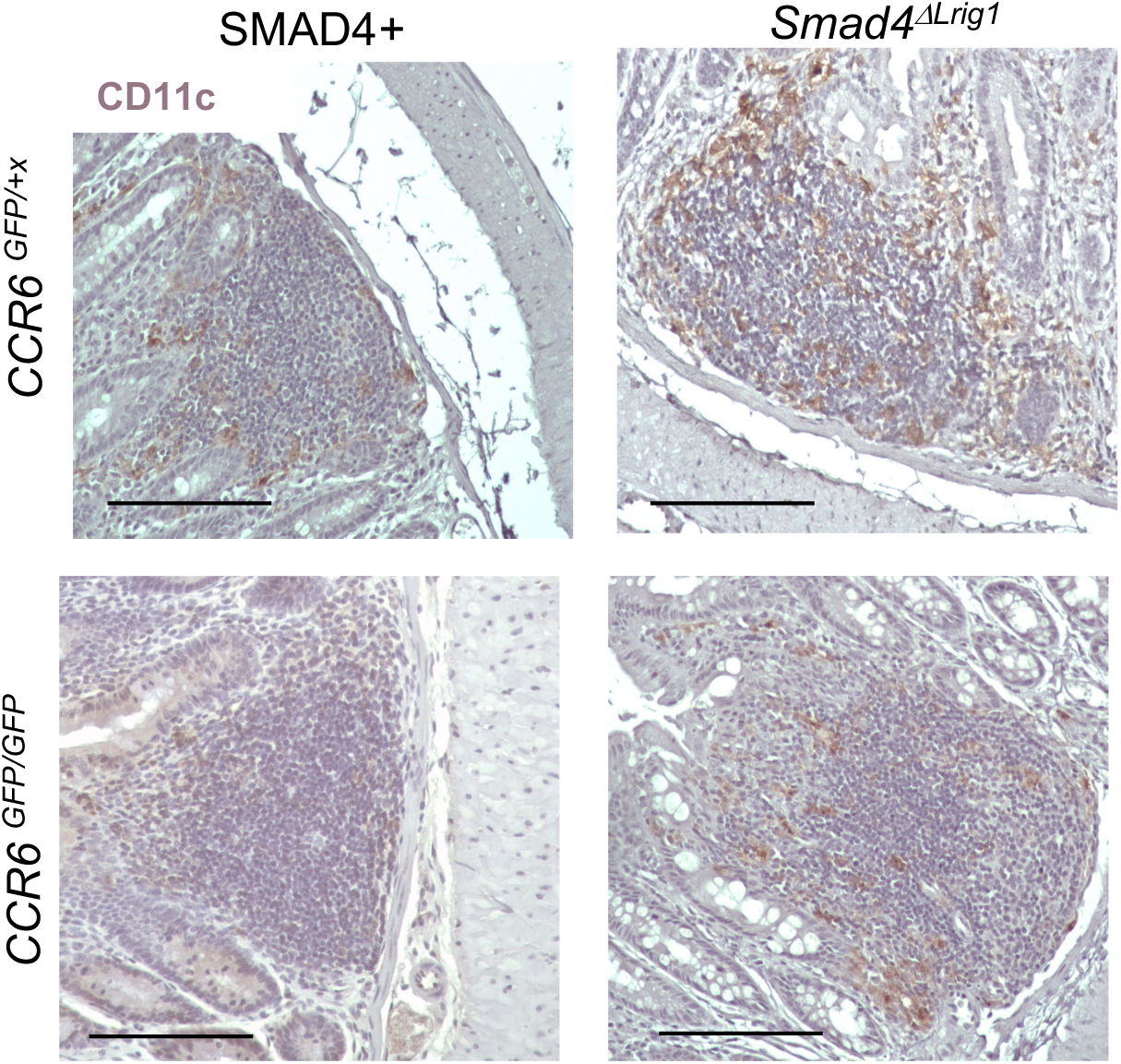
*Smad4* loss does not alter the distribution of CD11c+ dendritic cells 3 months after completion of DSS-induced colitis. Colonic tissue sections were stained for CD11c (brown) and counterstained with hematoxylin (blue) (scale bars: 100μm).

## References

1. Elinav E, Nowarski R, Thaiss CA, et al. Inflammation-induced cancer: crosstalk between tumours, immune cells and microorganisms. Nat Rev Cancer 2013;13:759–71.

2. Siegel RL, Miller KD, Goding Sauer A, et al. Colorectal cancer statistics, 2020. CA Cancer J Clin 2020;70:145–164.

3. Coussens LM, Werb Z. Inflammation and cancer. Nature 2002;420:860–7.

4. Smith PM, Means AL, Beauchamp RD. Immunomodulatory Effects of TGF-beta Family Signaling within Intestinal Epithelial Cells and Carcinomas. Gastrointest Disord (Basel) 2019;1:290–300.

5. Means AL, Freeman TJ, Zhu J, et al. Epithelial Smad4 Deletion Up-Regulates Inflammation and Promotes Inflammation-Associated Cancer. Cell Mol Gastroenterol Hepatol 2018;6:257–276.

6. Comerford I, Bunting M, Fenix K, et al. An immune paradox: how can the same chemokine axis regulate both immune tolerance and activation?: CCR6/CCL20: a chemokine axis balancing immunological tolerance and inflammation in autoimmune disease. Bioessays 2010;32:1067–76.

7. Wang L, Liu Q, Sun Q, et al. TLR4 signaling in cancer cells promotes chemoattraction of immature dendritic cells via autocrine CCL20. Biochem Biophys Res Commun 2008;366:852–6.

8. Frick VO, Rubie C, Kolsch K, et al. CCR6/CCL20 chemokine expression profile in distinct colorectal malignancies. Scand J Immunol 2013;78:298–305.

9. Skovdahl HK, Granlund A, Ostvik AE, et al. Expression of CCL20 and Its Corresponding Receptor CCR6 Is Enhanced in Active Inflammatory Bowel Disease, and TLR3 Mediates CCL20 Expression in Colonic Epithelial Cells. PLoS One 2015;10:e0141710.

10. Zhang H, Zhong W, Zhou G, et al. Expression of chemokine CCL20 in ulcerative colitis. Mol Med Rep 2012;6:1255–60.

11. Vancamelbeke M, Vanuytsel T, Farre R, et al. Genetic and Transcriptomic Bases of Intestinal Epithelial Barrier Dysfunction in Inflammatory Bowel Disease. Inflamm Bowel Dis 2017;23:1718–1729.

12. Marincola Smith P, Choksi YA, Markham NO, et al. Colon epithelial cell TGFbeta signaling modulates the expression of tight junction proteins and barrier function in mice. Am J Physiol Gastrointest Liver Physiol 2021;320:G936–G957.

13. Lugering A, Ross M, Sieker M, et al. CCR6 identifies lymphoid tissue inducer cells within cryptopatches. Clin Exp Immunol 2010;160:440–9.

14. Edgar R, Domrachev M, Lash AE. Gene Expression Omnibus: NCBI gene expression and hybridization array data repository. Nucleic Acids Res 2002;30:207–10.

15. Barrett T, Wilhite SE, Ledoux P, et al. NCBI GEO: archive for functional genomics data sets--update. Nucleic Acids Res 2013;41:D991–5.

16. Means AL, Ray KC, Singh AB, et al. Overexpression of heparin-binding EGF-like growth factor in mouse pancreas results in fibrosis and epithelial metaplasia. Gastroenterology 2003;124:1020–36.

17. Dieleman LA, Palmen MJ, Akol H, et al. Chronic experimental colitis induced by dextran sulphate sodium (DSS) is characterized by Th1 and Th2 cytokines. Clin Exp Immunol 1998;114:385–91.

18. Banerjee A, Herring CA, Chen B, et al. Succinate Produced by Intestinal Microbes Promotes Specification of Tuft Cells to Suppress Ileal Inflammation. Gastroenterology 2020;159:2101–2115 e5.

19. Habtezion A, Nguyen LP, Hadeiba H, et al. Leukocyte Trafficking to the Small Intestine and Colon. Gastroenterology 2016;150:340–54.

20. Rhee KJ, Jasper PJ, Sethupathi P, et al. Positive selection of the peripheral B cell repertoire in gut-associated lymphoid tissues. J Exp Med 2005;201:55–62.

21. Morbe UM, Jorgensen PB, Fenton TM, et al. Human gut-associated lymphoid tissues (GALT); diversity, structure, and function. Mucosal Immunol 2021;14:793–802.

22. Kucharzik T, Hudson JT, 3rd, Waikel RL, et al. CCR6 expression distinguishes mouse myeloid and lymphoid dendritic cell subsets: demonstration using a CCR6 EGFP knock-in mouse. Eur J Immunol 2002;32:104–12.

23. Varona R, Villares R, Carramolino L, et al. CCR6-deficient mice have impaired leukocyte homeostasis and altered contact hypersensitivity and delayed-type hypersensitivity responses. J Clin Invest 2001;107:R37–45.

24. Okayasu I, Yamada M, Mikami T, et al. Dysplasia and carcinoma development in a repeated dextran sulfate sodium-induced colitis model. J Gastroenterol Hepatol 2002;17:1078–83.

25. Gough NR, Xiang X, Mishra L. TGF-beta Signaling in Liver, Pancreas, and Gastrointestinal Diseases and Cancer. Gastroenterology 2021;161:434–452 e15.

26. Ikushima H, Miyazono K. TGFbeta signalling: a complex web in cancer progression. Nat Rev Cancer 2010;10:415–24.

27. Principe DR, Doll JA, Bauer J, et al. TGF-beta: duality of function between tumor prevention and carcinogenesis. J Natl Cancer Inst 2014;106:djt369.

28. Roberts AB, Wakefield LM. The two faces of transforming growth factor beta in carcinogenesis. Proc Natl Acad Sci U S A 2003;100:8621–3.

29. Yang L, Pang Y, Moses HL. TGF-beta and immune cells: an important regulatory axis in the tumor microenvironment and progression. Trends Immunol 2010;31:220–7.

30. Travis MA, Sheppard D. TGF-beta activation and function in immunity. Annu Rev Immunol 2014;32:51–82.

31. Massague J. TGF-beta signaling in development and disease. FEBS Lett 2012;586:1833.

32. Monteleone G, Kumberova A, Croft NM, et al. Blocking Smad7 restores TGF-beta1 signaling in chronic inflammatory bowel disease. J Clin Invest 2001;108:601–9.

33. Williams IR. CCR6 and CCL20: partners in intestinal immunity and lymphorganogenesis. Ann N Y Acad Sci 2006;1072:52–61.

34. Wunderlich CM, Ackermann PJ, Ostermann AL, et al. Obesity exacerbates colitis-associated cancer via IL-6-regulated macrophage polarisation and CCL-20/CCR-6-mediated lymphocyte recruitment. Nat Commun 2018;9:1646.

35. Li L, Boussiotis VA. The role of IL-17-producing Foxp3+ CD4+ T cells in inflammatory bowel disease and colon cancer. Clin Immunol 2013;148:246–53.

36. Ghazalsofala R, Rezaee SA, Rafatpanah H, et al. Evaluation of CD4+ CD25+ FoxP3+ Regulatory T cells and FoxP3 and CTLA-4 gene Expression in Patients wwith Newly Diagnosed Tuberculosis in Northeast of Iran. Jundishapur J Microbiol 2015;8:e17726.

37. Maul J, Loddenkemper C, Mundt P, et al. Peripheral and intestinal regulatory CD4+ CD25(high) T cells in inflammatory bowel disease. Gastroenterology 2005;128:1868–78.

38. Wu S, Rhee KJ, Albesiano E, et al. A human colonic commensal promotes colon tumorigenesis via activation of T helper type 17 T cell responses. Nat Med 2009;15:1016–22.

39. Bernardo D, Marin AC, Fernandez-Tome S, et al. Human intestinal pro-inflammatory CD11c(high)CCR2(+)CX3CR1(+) macrophages, but not their tolerogenic CD11c(-)CCR2(-)CX3CR1(-) counterparts, are expanded in inflammatory bowel disease. Mucosal Immunol 2018;11:1114–1126.

40. Gu T, Li Q, Egilmez NK. IFNbeta-producing CX3CR1(+) macrophages promote T-regulatory cell expansion and tumor growth in the APC(min/+) / Bacteroides fragilis colon cancer model. Oncoimmunology 2019;8:e1665975.

