## Supplementary Methods for "SMAD4 suppresses colitis-associated carcinoma through inhibition of CCL20/CCR6-mediated inflammation"

*In silico Analysis of Microarray Data and The Cancer Genome Atlas (TCGA)*

Transcriptomic data from colonoscopic biopsy specimens of IBD patients and healthy controls (data accessible at NCBI GEO database, accession number GSE75214) included colon mucosal biopsies from the following: 74 patients with active UC (UCa), 23 patients with inactive UC (UCi), 8 patients with active Crohn’s Disease (CDa), and 11 healthy controls (HC).^1^ mRNA-seq expression data from the TCGA database included the following: 82 primary colon cancer (CC) specimens, 41 healthy control colon (HCc) specimens, 99 primary rectal cancer specimens (RC), and 10 healthy rectum control (HCr) specimens.

*Multiplex Immunofluorescent (mxIF) staining of Human Tissues*

MxIF imaging was performed on an Olympus X81 inverted microscope with filter sets specific for DAPI, Cy2, Cy3, and Cy5 in the VUMC DHSR. TMAs were stained with antibodies against β-Catenin, NaK-ATPase, panCK, SMA, vimentim, PCNA, γ-actin, CD3, CD4, CD8, CD11b, CD20, CD68, and FoxP3.

*RNA-Sequencing*

Following crypt chelation, the sub-epithelial stroma from three *Smad4^ΔLrig1^* and three SMAD4+ control mice was removed from the underlying muscle layer by scraping. RNA was then made from the sub-epithelial stroma using RNeasy kit (Qiagen, Germantown, MD) and RNAseq performed by the Vanderbilt Technologies for Advanced Genomics (VANTAGE) core facility. 32–37 million 51–base pair, single-end reads were generated per sample. Reads were mapped to the mouse genome mm10 using TopHat-2.1.0, uniquely mapping 86%–95% single-end reads to the genome. The number of reads that fell into annotated genes were counted using samtools-1.3.1 and HTSeq-0.5.4p5. Count-based differential expression analysis was performed using edgeR_3.4.2.

Pathways and upstream regulators were analyzed using Ingenuity Pathway Analysis (IPA) software (Qiagen, Inc, <https://www.qiagenbioinformatics.com/products/ingenuity-pathway-analysis>).^2^ Immune cell composition was predicted using ImmuCC software (<http://218.4.234.74:3200/immune/>) as previously described.^3^

*RNAScope ®* In Situ *Hybridization*

*In situ* hybridization by RNAscope® technique was performed according to the manufacturer’s instructions [Advanced Cell Diagnostics (ACD), Newark, CA, USA]. Briefly, 5μm thick sections were deparaffinized and treated with hydrogen peroxidase followed by antigen retrieval and protease treatment. Probes for *Ccl20* and *Ccr6*, as well as negative and positive controls provided by the manufacturer, were used. The probes were then hybridized for 2 hours followed by 10 amplification steps. The signal was detected by RNAscope® 2.5 HD Duplex Reagent Kit and visualized via brightfield microscopy. To quantify differential *Ccl20* expression, mRNA dots were counted in each positive cell. To quantify *Ccr6* expression, the number of positive cells were counted.

*Histology and Immunostaining*

For immunohistochemistry (IHC) and immunofluorescence (IF), the following antibodies were used: rabbit anti-GFP (NB600-308, Novus Biologicals), rat anti-B220 (BD-553084 BD Biosciences, San Jose, CA), rabbit anti-CD11c (97585S, Cell Signaling Technology, Danvers, MA), goat anti-CD3 (SC-1127, Santa Cruz Biotechnology, Santa Cruz, CA), anti-FoxP3 (Ab54501, Abcam, Cambridge, MA), and anti-mouse IL17 (SC-37428, Santa Cruz Biotechnology). Cy2-, Cy3-, and Cy5- conjugated secondary antibodies used in IF were purchased from Jackson ImmunoResearch Laboratories (West Grove, PA).

For immunohistochemistry (IHC), brightfield images were captured on an Axioskop 40 microscope using Axiovision software (Carl Zeiss Microimaging, Thornwood, NY). Fluorescent images were captured with the Zeiss LSM 510 Meta Inverted Confocal Microscope (Carl Zeiss Microimaging) through the Cell Imaging Shared Resource at VUMC. IHC and IF images were processed with ImageJ processing software (National Institutes of Health, Bethesda, MD) and cell quantification was performed using the ImageJ cell counter feature.

*Flow Cytometry*

Epithelial and stromal cell suspensions were created from the colons of *SMAD*4+*Ccr6 ^GFP/+^,* SMAD4+*Ccr6^GFP/GFP^, Smad4^ΔLrig1^Ccr6^GFP/+^*, and *Smad4^ΔLrig1^Ccr6 ^GFP/GFP^* mice. Both epithelial and stromal cell suspensions were treated with a commercially available Cell Stimulation Cocktail (Invitrogen) per manufacturer’s instruction. Treated cell suspensions were then incubated for seven hours at 37°C and washed prior to further staining. Cells were stained with a Live/Dead™ fixable viability dye (Invitrogen) prior to surface and intracellular staining to distinguish live cells. Single cell suspensions from both the epithelium and stroma were aliquoted for staining in two parallel panels. Antibodies for the lymphoid and myeloid panel are displayed in Supplementary Figure 1a (BioLegend, San Diego, CA). The anti-GFP antibody was titrated to stain specifically cells that normally express CCR6. Thus, cells stained for GFP+ were also CCR6+ (Supplementary Figure 1b). The cell suspensions were first stained for surface markers, incubated on ice for 1 hour in the dark and resuspended in FACS buffer prior to fixation and permeabilization using Intracellular Fixation & Permeabilization buffer set (Invitrogen) as directed. Cells were then stained for intracellular markers, which included antibodies against FoxP3, IL-17a, and GFP. The cell suspensions were then washed and resuspended in FACS buffer prior to FACS analysis. Gating and analyses were performed using FlowJo 10 software (FlowJo, Ashland, OR).

References

1.	Vancamelbeke M, Vanuytsel T, Farre R, et al. Genetic and Transcriptomic Bases of Intestinal Epithelial Barrier Dysfunction in Inflammatory Bowel Disease. Inflamm Bowel Dis 2017;23:1718-1729.

2.	Kramer A, Green J, Pollard J, Jr., et al. Causal analysis approaches in Ingenuity Pathway Analysis. Bioinformatics 2014;30:523-30.

3.	Chen Z, Quan L, Huang A, et al. seq-ImmuCC: Cell-Centric View of Tissue Transcriptome Measuring Cellular Compositions of Immune Microenvironment From Mouse RNA-Seq Data. Front Immunol 2018;9:1286.
